# Genomic landscape of antiviral defense systems in prokaryotes and phages

**DOI:** 10.1101/2024.10.26.620412

**Authors:** Jinquan Li, Jiazheng Gu, Runyue Xia, Meng Li

**Author notes:** These authors contributed equally to this work (Jinquan Li, Jiazheng Gu).

## Abstract

Prokaryotes and their viruses have co-evolved for billions of years, resulting in emergence of numerous antiviral defense systems. With the development of bioinformatic technologies and experimental studies, more and more novel defense systems have been discovered in the near decades. However, the origin and mechanism of these systems are still largely unknown. This study provides a systematic analysis of 132 defense systems within 212,599 prokaryotic genomes, which should be the largest analyzed data so far, revealing the diversity and distribution of these systems across different taxonomic units. Our findings also reveal that not only well-studied bacteria, but also archaea and virus encode diverse antiviral defense systems with specific features. In summary, this work classify 132 known defense systems, provides a comprehensive view of prokaryotic defense systems and insights into the evolution of immune responses.

## 1. Introduction

Bacteriophages (phages) are viruses that infect microorganisms and are one of the most abundant life forms on Earth [1]. The widespread presence of prophage and proviral sequences in bacterial and archaeal genomes demonstrates that most prokaryotes have been infected by their viruses [2]. Current research suggests that virus evolved billions of years ago, shortly after the emergence of prokaryotes. It has been hypothesized that the arms race between prokaryotes and their virus has been going continuously during the evolution [3]. A dynamic equilibrium is presents between the competition of prokaryotes and their viruses in natural conditions, and these interactions can have diverse consequences. For example, prokaryotes are invaded by a wide range of mobile genetic elements (MGEs) such as phages, plasmids and transposons that confer new phenotypes to the host, such as resistance to antibiotics [4,5]. Alternatively, phages can affect the genetic stability of the host, and often lead to the death of the infected host cell. Therefore, prokaryotes possess defenses against genetic invasion, and MGEs possess a set of mechanisms for evading these defenses [6,7].

Antiviral defense systems in prokaryotes are similar to the immune systems of eukaryotes, and can be divided into two main categories, innate and adaptive. Restriction modification (RM) [8] and clustered regularly interspaced short palindromic repeat (CRISPR)-CRISPR-associated (Cas) [9] systems represent examples of these two types of immune systems, respectively, and they are also the two most widely distributed and well-studied defense mechanisms. RM systems are present in >90% of sequenced prokaryotic genomes [10], whereas CRISPR-Cas systems are present in about 30-40% of bacterial and >90% of archaeal genomes [11].

Antiviral defenses can result in a process known as the abortive infection (Abi) and is considered a defense of last resort. Defenses that function via Abi prevent phages from completing their life cycle by causing the infected cell to undergo self-sacrifice, which protects the host population through reducing the spread of the viral epidemic. These systems are normally in an “off-state” until activated by recognition of proteins or nucleic acids of the invading phages or changes in the intracellular physiology caused by infection [11,12]. Toxin-antitoxin (TA) systems are a large class of Abi systems consisting of toxin and antitoxin (proteins or RNA). The toxins can inhibit bacterial growth or even kill the host, while antitoxins neutralize the toxin. TAs typically maintain a balance with no effect on normal bacterial growth until phage infection disrupts that balance and elicits Abi [13].

Genes encoding defense system often occur in clusters called “defense islands” [14,15]. To date, greater than 100 defense systems have been validated [16]. However, these varied systems look sophisticated and their distribution across different taxonomic units is less studied. Here, we systematically reviewed and analyzed the distribution of 132 defense systems within 212,599 genomes using DefenseFinder [17] (August 2023), and classified them according to the structural domains of their putative effector proteins. Moreover, we calculated the distribution and occurrence frequency of these systems within different prokaryotic taxonomic units, bringing some insights into the ecological and evolutionary situation of defense systems.

## 2. Classification of defense systems based on functional domains

Currently known defenses are so diverse that the mechanisms and relationships between them remain poorly understood. Simply, based on the number of genes that make up the defense systems, they can be classified as monogenic and polygenic systems. For these 132 defense systems, 61 systems comprise single genes (e.g. AbiA, SEFIR, Shedu), 49 systems encode two genes, 16 systems encode three genes, whereas 6 systems encode four genes or more. Here, we first classified these defense systems primarily based on their functional domains (Figure 1), which could reflect their potential mechanisms. The defense systems are mainly assembled by sensing domain and effector domain modules, among them different combinations result in different systems. To comprehensively organize these defense systems, we distinguished them primarily according to the differences of enzyme activities of their effectors.

**Figure 1.**
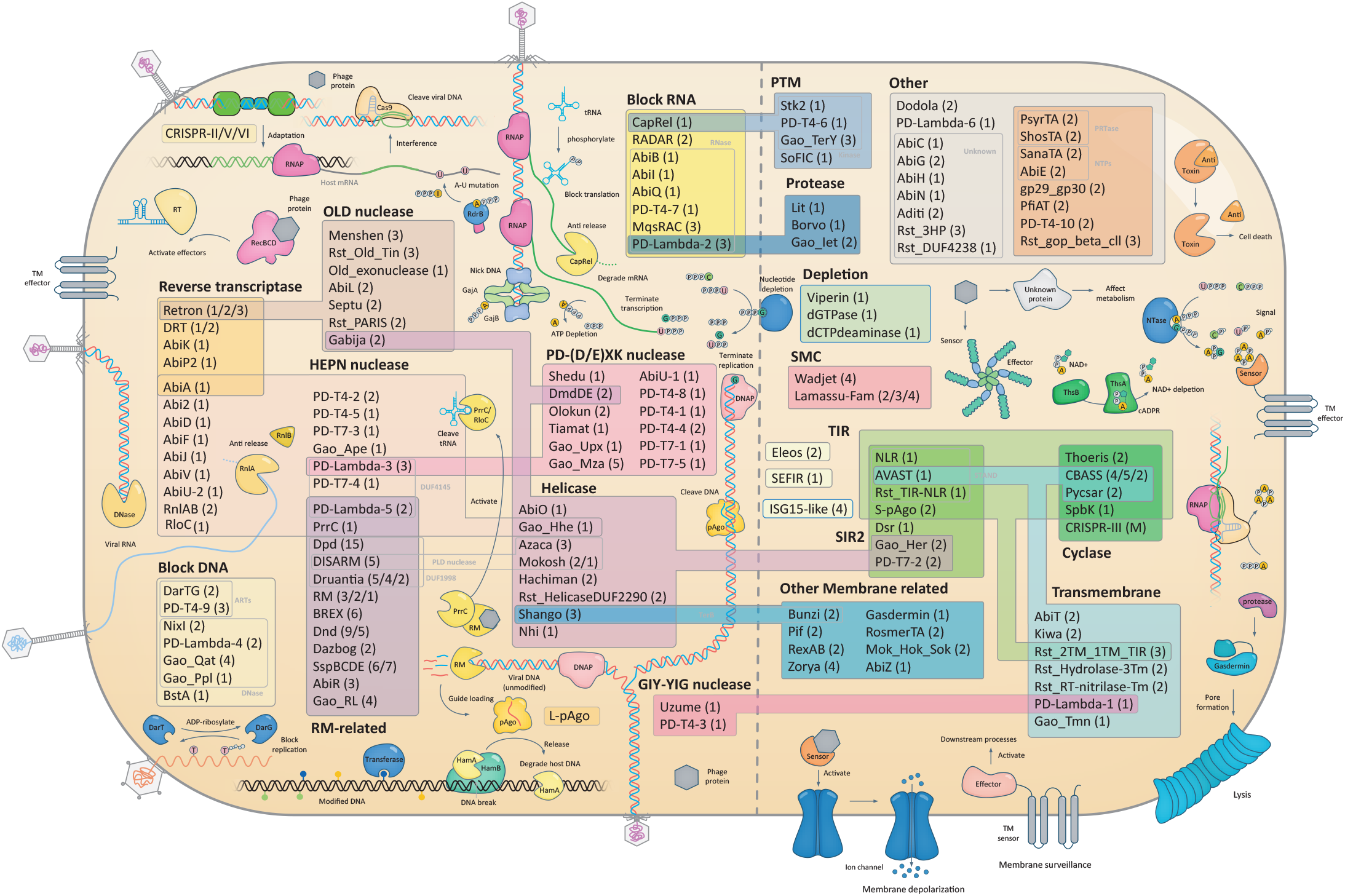
**Classification of 132 defense systems based on their functional structural domains and mechanism**. Defense systems consist of HEPN domain-containing protein are clustered with flesh color background. In AbiA system, a reverse transcriptase domain is fused to the HEPN domain. Five systems consisting of reverse transcriptase domain are in orange background color. Twelve systems consisting of proteins that can modify DNA or distinguish the modified DNA are thought to be related with RM system, are colored in purple background. Defense systems consist of helicase domain-containing protein are in pink background. Defense systems consist of PD-(D/E)XK nuclease domain-containing protein are in khaki color. Defense systems consist of OLD nuclease domain-containing protein are in cameo color. Seven defense systems with diverse domains can block DNA replication of virus are colored in silver gray. Eight defense systems with diverse domains can block RNA transcription or translation are colored in gold color. CapRel function by pyrophosphorylating the tRNA, thus also clustering with other 4 systems that involve in post-translational modification (PTM). PD-Lambda-2 system consists of protein with peptidase domain, therefore clusters with 3 other systems related with protease. Defense systems associated with TIR domain are colored green, while blackish green for SIR domain, olive green for cyclase domain. Defense systems that function by depleting the nucleotides are in mint green. Defense systems consist of transmembrane domain protein are in cyan color, other systems associated with membrane are in sky blue. Eleos, SEFIR, and ISG15-like systems, which are related with eukaryotic immunity, are in cream color. Defense systems associated with SMC proteins are in magenta color. Other systems that cannot be classified in a clear group are colored gray, while TA systems are further clustered with brown color.

### 2.1 Nucleic acids related systems

Most defense systems act directly on the nucleic acids of viruses to inhibit their propagation, or on nucleic acids of the host microbe to arrest growth. Therefore, we first classified a group of systems which function through nucleic acids manipulation. These systems consist of a sensor domain that sense invading nucleic acids, or an effector domain that cleaves DNA or RNA of the virus or host.

#### 2.1.1 HEPN nuclease domain

The HEPN (Higher Eukaryotes and Prokaryotes Nucleotide-binding) domains are widespread in defense systems and were initially thought to be associated with DNA polymerase β-type nucleotidyltransferase in prokaryotes and sacsin proteins in animal [18,19]. In eukaryotes, HEPN domains are involved in RNA processing, stress responses and innate immunity [20–23]. In prokaryotic defenses, most HEPN domain-containing proteins have RNase activity which inactivates viral RNA by cleavage, or mediates Abi by cleaving host nucleic acids. RloC and PrrC are anticodon nucleases (ACNases) that cleave tRNA of the host to inhibit viral protein synthesis [24,25]. The HEPN superfamily is also found within some related type II TA systems, where HEPN activity is mitigated by the antitoxin minimal nucleotidyltransferase (MNT) domain-containing protein [26,27]. For example, the RnlA toxin in the RnlAB system is an RNase that directly attacks the viral nucleic acids in most cases [28].

HEPN domains are present in 18 systems, including 7 Abi systems (Abi2, AbiD, AbiF, AbiJ, AbiU-2, AbiV, C-terminus of AbiA) originally found in *Lactococcus lactis*, 7 single-gene systems (PD-T4-5, PD-T7- 3, PD-T7-4, Gao_Ape, Gao_Hhe, PrrC, RloC) and 4 multiple-gene systems (RnlAB, PD-T4-2, PD-Lambda- 3, PD-Lambda-5). HEPN domain-containing proteins are also present in type I and III CRISPR-Cas systems, including Cas13, Csx1, and Csm6 [29]. In brief, HEPN domain has RNase activity in most cases, and participates in many metabolic activities to process RNA. Therefore, they can act as effectors in numerous antiviral defense system to cleave the viral or host RNA.

#### 2.1.2 OLD family nuclease domain

The OLD (Overcoming Lysogenization Defect) family protein was firstly discovered in P2 phage [30], which is an ATP-powered nucleases, consisting of an N-terminal ATPase domain and a C-terminal Toprim (Topoisomerase-primase) domain. The toxicity of OLD proteins requires both the ATPase and Toprim domains [31,32]. OLD family proteins enable P2 to repel the infection of other phages [30], and are also present within 8 defense systems: Gabija, PARIS, Septu, Menshen, AbiL, Old_exonuclease, Rst_Old_Tin, type I-B1/B2 Retron. Three of the best-characterized systems are discussed below.

The Gabija system consists of two genes, GajA and GajB [33] and the structure revealed that GajA contains both ATPase and Toprim domain characteristic of OLD family. GajA has a region for dimerization and assembly [34,35], which acts as a sequence specific endonuclease in the Gabija system. Its activity is strictly regulated by the nucleotide concentration, and it will be activated only when NTP and dNTP are consumed in large quantities. GajB activates GajA to media effective abortion infection defense against phages by regulating nucleotide hydrolysis [36].

PARIS system is consisting of two genes, AriA and AriB, where AriA is an ATPase, while AriB contains DUF4435 protein with weak homology to the Toprim domains [37]. Recent study showed that PARIS is activated by the Ocr protein of phage [38], which has been shown to resist RM and BREX defense systems [39,40]. The PARIS system is therefore considered to be resembled to OLD family defense systems.

Septu system also consists of two genes, PtuA and PtuB [33], where PtuA contains an ATPase domain related with OLD proteins, while PtuB is a putative HNH family nuclease. This system can exist independently as a stand-alone system or be associated with retrons [41–43]. Structural and functional analysis showed that PtuA and PtuB assemble into an inflammasome-like complex with a stoichiometry of 6:2, functioned by cleaving phage genomic DNA and regulating by ATP molecule [44].

#### 2.1.3 PD-(D/E)XK superfamily nuclease domain

The PD-(D/E)XK superfamily is a group of magnesium-dependent nuclease. It was initially identified in type II restriction endonuclease (REase) [45], and later in many proteins that fill in a variety of biological functions, such as DNA recombination, DNA repair, tRNA splicing [46–49]. This superfamily is characterized by the presence of a central 4 strands β-sheet, flanked by two α-helices on both sides (αβββαβ topology) [50]. Previous studies have found many unknown proteins with PD-(D/E)XK structure using sequence alignment and HHsearch methods [51], suggesting that PD-(D/E)XK superfamily proteins are widely distributed and related with nucleic acid metabolism [52].

Our category shows that 13 defense systems contain PD-(D/E)XK structure in their coding genes. Among them, 8 systems (Shedu, Tiamat, Gao_Upx, AbiU-1, PD-T4-8, PD-T4-1 PD-T7-1, PD-T7-5) consist of single gene, suggesting these systems consist of a nuclease, and therefore may act directly on the nucleic acids.

#### 2.1.4 GIY-YIG nuclease domain

The GIY-YIG nuclease superfamily shares a common core fold, comprising of about 100 aa domain with motif GIY-YIG in the N-terminal, followed by an Arg residue in the center and a Glu residue in the C-terminal [53]. Similar to the PD-(D/E)XK family nuclease, GIY-YIG nuclease is also an endonuclease involved in the destruction and reconstruction of phosphodiester bonds. It was originally discovered in homing endonuclease (HEase), and later identified in many proteins involved in genome degradation, DNA repair, recombination, and transpose [54–56].

Unlike the PD-(D/E)XK family nuclease, GIY-YIG nuclease has a strictly conserved active site, and its six key catalytic sites are strictly conserved. This led to the relatively stable and single evolution of GIY-YIG nuclease [57]. In view of the ability of GIY-YIG nuclease to cleave nucleic acids, a domain homology to GIY-YIG family is detected in 3 single-gene systems (PD-T4-3, PD-lambda-1, Uzume), probably involved in nucleic acids degradation.

#### 2.1.5 Reverse transcriptase

Reverse transcriptases (RTs) catalyze the polymerization of DNA from an RNA template. Traditionally, most RTs are components of mobile retroelements in prokaryotes [58]. Retrons are one of the genetic elements consisting of RTs and non-coding RNA. The Retron system is composed of RT, non-coding RNA, and effector protein, where the effector proteins can be a different combination of those structural domains, such as nucleases, or proteases, etc. [42]. Abi systems of AbiA, AbiK, and AbiP2 systems are also related with RT domain [59–61].

In 2020, six groups of previously unknown RTs which lack the features of mobility were showed to have antiviral activities, named defense associated RTs (DRTs) [62]. Among them, a variety of DRTs show different phase resistance. DRT type 1 is thought to suppress the expression of phase late genes. DTR type3 does not directly inhibit phage gene expression, and structured noncoding RNA may have an important impact on its activation [62].

#### 2.1.6 Helicase domains

Helicases normally act on the DNA to unwind the complementary strands [63]. The defense system that encodes helicase domains is usually composed of several genes that work together to play an anti-phage role. For example, Hachiman system is a two gene cassette consisting of HamA, the effector nuclease, and HamB, the sensor helicase [33]. Recent structural and functional analysis showed that it forms heterodimeric nuclease-helicase complex, where HamA is constrained by HamB binding to intact DNA. When the system detects DNA damage, HamA is released from the complex, degrading all the DNA of both phage and host [64].

Similarly, we hypothesize that other systems with nuclease-helicase architecture have similar mechanisms of function. Rst_HelicaseDUF2290 system was shown to resist T7 phage, and its structure consists of an ATP dependent helicase connected with a DUF2290 protein [37]. Azaca system consists of helicase, PLD (phospholipase-D) nuclease, and DUF6361 [65]. The Mokosh system which consists of RNA helicase and PLD nuclease was thought to recognize phage RNA or RNA/DNA intermediates associated with phage replication [65]. AbiO, DmdDE, Shango, Nhi, Gao_Hhe, Gao_Her, and PD-T7-2 systems also encode a helicase.

#### 2.1.7 RM related

The RM related systems function by recognizing specific sequences on bases to distinguish between self and foreign genetic element. The well-established RM system can be classified into I-IV four subtypes based on their compositions and mechanisms [8,66]. In type I-III systems, host gene sequences are usually methylated by methyltransferases *in vivo*, and modified host genes cannot be degraded by restriction endonucleases. Methyltransferases are not present in the type IV system, so this system targets modifications on foreign genes for their cleavage [67].

Several other systems have similar features. For example, BREX, Dnd, SspBCDE, DISARM, Dpd, Druantia, PD-Lambda-5 and Gao_RL systems encode proteins that can modify DNA, while Dazbog, PrrC, and AbiR systems were showed to sense modified DNA [68]. Therefore, these systems are thought to be functioned by the way closely associated with RM systems. Here, we summarized some well-established systems in detail.

For BREX system, the most widespread type I BREX system employs an RM like principle, which its BrxX methylates the host DNA at specific sites, so the unmethylated phage DNA cannot replicate [69]. In addition to this, the BREX system is often combined with SAM (S-adenosyl-methionine) to enhance DNA binding [70–72]. Unlike using methyl as a marker to distinguish between self and non-self nucleic acids, the Dnd system confers nuclease resistance by phosphorothioate (PT) modification, which replaces the non-bridging oxygen on the DNA sugar-phosphate backbone with sulfur [73,74]. Beyond that, the Dnd system can enhance defense function by working together with some other system like the SspBCDE system [74]. SspBCDE system produces PT modifications on host genomic DNA, and SspE acts as a restriction part capable of sensing PT modifications on DNA to distinguish between the host and foreign genetic elements [75,76]. The DISARM system comprises four genes, including a DNA methylase and three other genes annotated as helicase domain (DrmA), DUF1998 domain (DrmB) and PLD nuclease domain (DrmC). DISARM methylase modifies host genomes as a marker of self nucleic acids similar to RM systems, but with different modified motifs [77,78]. The Dpd system utilizes multiple enzymes (DpdA-K) to substitute guanine residues with 7-deazaguanine derivatives. Interestingly, DpdB, like DndB in the Dnd system, has a role in regulating the expression of the Dpd modification genes [3,79]. The defense mechanism of the Druantia system is still not well understood, but its antiphage activity can be counteracted by hmC modification, indicating that this system may utilize hmC in host genomes as a defense marker similar to the RM system. It has been demonstrated that prrC is genomically linked to prrI RM system, whose three core components (hsdMSR) negatively regulate the PrrC by keeping it in a catalytically inactive state [80].

#### 2.1.8 Other nucleic acids related systems

Structural maintenance of chromosome (SMC) protein, which is involved in the maintenance and stabilization of chromosome structure in eukaryotes [81], also found to be related with prokaryotic defenses in Lamassu-Fam and Wadjet systems [33,65]. Lamassu-Fam systems mainly comprise two core proteins, where LumA constitutes the effector of the systems, and LmuB contains a SMC domain predicted to bind DNA [65]. Recent studies revealed some evidence that these systems with SMC protein recognize invasive linear or circular DNA [82]. The SMC family protein in the Wadjet system, JetC, can activate the nuclease subunit JetD specifically to cleave the plasmid DNA by sensing the topology of the circular DNA in the presence of ATP [83,84].

We further classified a group of systems function by blocking the DNA replication of viruses, which mainly include 7 systems (DarTG, PD-T4-9, BstA, NixI, PD-Lambda-4, Gao_Qat, Gao_Ppl). DarTG and PD-T4-9 TA systems encode ADP-ribosyltransferases (ARTs), where DarT toxin is an ART that modifies thymidines on single-stranded DNA in a sequence-specific manner [85,86]. BstA with a putative DNA-binding domain was shown to prevent phage DNA replication [87]. NixI, PD-Lambda-4, Gao_Qat, and Gao_Ppl systems consist of DNase that may degrade viral or host DNA directly.

We also classified a group of systems function by blocking the RNA transcription or translation of viruses, which mainly include 8 systems (CapRel, RADAR, PD-Lambda-2, MqsRAC, AbiB, AbiI, AbiQ, PD-T4-7). For CapRel system, it is a fused TA system which N-terminal nucleotide pyrophosphokinase toxin is rescued by the C-terminal antitoxin domain. Upon phage infection, the major capsid protein binds directly to the C-terminal structural domain to relieve autoinhibition, enabling the toxin to pyrophosphorylate the tRNA, thereby inhibiting translation [88]. RADAR (phage restriction by an adenosine deaminase acting on RNA) system is consists of an ATPase and a divergent adenosine deaminase. This system edits RNA transcripts by catalyzing the deamination of A to G, blocking the early genes of the phage [62]. PD-Lambda-2 system encodes a HigB toxin, which previously found to be associated with 50S ribosomal subunit, by cleaving within mRNA coding regions with AAA sequences [89]. In MqsRAC system which consists of a conventional type II TA and a molecular chaperone, MqsR serves as a RNase toxin that cleaves mRNA at GCU sites [90]. Other 4 systems (AbiB, AbiI, AbiQ, PD-T4-7) also express RNase activity, although the mechanisms still remain to be elucidated.

### 2.2 Membrane related systems

An important strategy for cells to defense against viruses is to protect the population by sacrificing the infected cells. This can be accomplished by lysing the cellular membrane by various processes, including cell membrane perforation, altering the membrane potential, etc. [91,92].

Protein elements that affect membrane permeability or integrity are widely found in the defense systems. They exist in organisms as a last line of defense. It is generally believed that this irreversible mode of defense is only selected in the late stages of defense. Therefore, membrane-related defense systems are usually inseparable from abortion and infection. Once membrane permeability is compromised, cell death is inevitable.

#### 2.2.1 Transmembrane domain

Transmembrane (TM) proteins are involved in many essential cell processes such as protein trafficking, energy conversion, signal transduction, and immune response [93–95]. The defining feature of transmembrane proteins is one or more TM domain, which are composed of α-helical structure with hydrophobic side chains or some β-sheet structure where they interact with the nonpolar lipid parts of the membrane [96,97]. On one hand, proteins with TM domain can lead to the destruction of membrane integrity, that ultimately lead to abortion infection. On the other, TM proteins can be the sensors to surveil the membrane integrality, and further activate downstream effectors.

For example, the cyclic oligonucleotides synthesized by CD-NTase in CBASS system can activate the TM proteins later, which are predicted to form pores in the membrane [98]. Pycsar system encodes a TM domain, which leads to loss of membrane integrity [99]. Some AVAST subtypes use TM domains as effectors to form pore, which are activated by sensing phage protein through the NLR domains [62,100]. It has also been reported that when the retron ec48 protein senses phage-induced RecBCD inhibition, it will activate the TM protein to cause host cell death [42]. KwaA protein of Kiwa system contains a TM domain, while KwaB acts as the effector to initiate a RecBCD-mediated process that reduces the efficiency of phage DNA replication [33,101]. Recent study by Rousset et al. found some diverse defense systems with TM domains enriched in phage hotspots, such as Rst_Hydrolase-3Tm, Rst_RT-nitrilase-Tm, and Rst_2TM_1TM_TIR [37]. Other systems like AbiT, PD-Lambda-1, Gao_Tmn also encode TM domain-containing proteins.

#### 2.2.2 Other diverse proteins acting on membrane

In addition to defense systems which directly consist of TM domain-containing proteins, there are other diverse proteins involved in defense related with cell membrane. We categorized 9 systems in this group as below.

Gasdermin is a family of recently identified pore-forming effector proteins that promote cytolysis and release cytokines to induce lytic cell death [102–104]. The formation of gasdermin pores is triggered by caspase mediated cleavage during inflammasome signaling, and is essential for defense against pathogens and cancer [105]. Recent study discovered prokaryotic homologue of gasdermin, which contains a conserved pore-forming domain, and is usually encoded nearby other protease [106]. Similar with eukaryotic gasdermin, bacterial gasdermin assembles into pore of uneven size after activation by cleavage of the inhibitory C-terminal peptide, and further disrupts membrane integrity to mediate cell death.

Zorya system was recently shown to be related with membrane [33], in which ZorA and ZorB form a transmembrane rotary motor heteromer with 5:2 stoichiometry. ZorA located inside the cytoplasm can further transfer the phage invasion signal to activate soluble ZorC and ZorD nuclease to prevent phage propagation [107]. For RexAB system, its RexB protein will assemble into ion channel to destroy the membrane permeability after the RexA protein senses viral DNA complex [108–110]. AbiZ is an Abi system that can prevent early transcription of phages [111]. Previous study showed that AbiZ is a protein anchored to the membrane by an N-terminal membrane spanning domain, and it induces cell death by damaging the cell membrane of the infected cell [111]. Similar with AbiZ system, Mok_Hok_Sok system was showed to target the inner membrane, and in turn triggers depolarization of the membrane [112,113]. For Pif system, activation of PifA toxicity by T7 capsid protein gp10 leads to leakage of ATP through loss of membrane integrity [114,115]. Both Bunzi and Shango system encode TerB domain-containing proteins which are associated with the periplasmic membrane, therefore they are considered to be the system related with membrane surveillance [65,116]. RosmerTA system consists of an unknow function toxin RmrT which can increase membrane depolarization and permeability, and an Zn peptidase antitoxin RmrA [65,117].

### 2.3 Signal transduction and eukaryotic immunity related systems

#### 2.3.1 TIR and Sir2 domain

Toll/Interleukin-1 receptor (TIR) domain and silent information regulator 2 (Sir2) were initially thought to be the important component of the immune response associated with signal transduction in plant and animal cells [118–120]. When the receptor comes into contact with virus, it will activate the TIR and Sir2 domains and produce a variety of cyclic ADP-ribose, which further leads to the death of the infected cells through signaling molecules [121–123].

This multicomponent coordinated defense process used to be considered unique for eukaryotes. However, as more and more prokaryotic defense systems being discovered, it has been found to be widespread from bacteria to archaea [124,125]. The cyclic di-GMP produced by the CBASS system [126], and the cyclic UMP produced by Pycsar system [99] have similar signals. TIR domain-containing proteins behave NADase activity that hydrolyzes NAD+ in these systems, thus trigger growth arrest through the depletion of cellular NAD+. Thoeris system is consists of ThsA (Sir2 domain) and ThsB (TIR domain), where TIR domain of ThsB acts as a sensor that generates an isomer of cyclic ADP ribose, which further activate ThsA to depletes NAD+ through its Sir2 domain [33,124].

TIR and Sir2 domains can also be involved as separate effector modules in many other defense systems. For example, TIR domains are associated with nucleotide cyclase in the CBASS and Pycsar systems [99,126], associated with reverse transcriptase in Gao_Retron-TIR system [62]. Some system subtypes of pAgo, Lamassu-Fam, Retron, etc. use these domains as effectors to deplete NAD+ as well [42,65,127]. For short pAgo system which contains only MID and PIWI domains, it typically associated with Sir2, Mrr or TIR domain-containing proteins as effectors [128]. NACHT-containing proteins are also typical proteins of defense systems, which comprise of three domains structure with a C-terminal sensor, a central NACHT and an N-terminal effector domain. Effector domain here usually carry TIR or Sir2 domains [129]. Other systems like Dsr, Gao_Her, PD-T7-2, AVAST V type, and Rst_TIR-NLR also encode TIR or Sir2 domains. The above examples illustrate that the TIR and Sir2 domains serve as the effector with NADase activity in diverse sets of antiviral defense systems.

#### 2.3.2 STAND superfamily

The STAND (signal transduction ATPases with numerous domains) superfamily is signaling protein including numerous NTPases that mediate programmed cell death and various forms of signal transduction in eukaryotes [130]. This superfamily, which typically contains a helical sensor domain and a effector domain, also has been found in bacteria and archaea [131]. For example, nucleotide-binding domain and leucine-rich repeat containing gene family (NLR), is characterized by NACHT module that belongs to STAND NTPases [129]. Recent studies showed that AVAST (Antiviral ATPases/NTPases of the STAND superfamily) and NLR-like systems detect virus and become oligomerization resemble inflammasome, further activate its effector domains like nucleases, TIR, or SIR, etc. to abrogate infection [100,129].

#### 2.3.3 Cyclase domain

Second messengers, including cyclic nucleotides (cAMP, GMP), Ca2+, NO, etc., play an important role in cellular signaling by rapidly activating cascade systems and regulating downstream responses [132–134].

The cyclic GMP-AMP synthase (cGAS)-STING pathway is a central component of the innate immune system in eukaryotes [135,136]. The cGAS recognizes viral DNA, and catalyze the generation of 2’3’- cGAMP from ATP and GTP. This molecular further acts as a second messenger to trigger the innate immune response mediated by the stimulator of interferon genes (STING) [137].

Antiviral singling pathways have recently been discovered in bacteria, where cGAS/DncV-like nucleotidyltransferases (CD-NTases) family proteins use purine and pyrimidine nucleotides to synthesize a diverse range of cyclic nucleotides to initiate antiviral response [138]. Previous studies in bacteria showed that the cyclic nucleotide-based antiphage signaling systems (CBASS) is activated by cGAS upon sensing phage infection [139]. The production of cGAMP as a second messenger further activates phospholipase effector proteins, leading to degradation of the cell membrane and host death [140]. The prokaryotic CBASS system is considered the ancestor of the eukaryotic cGAS-STING signaling pathway. Another newly discovered defense system involved in signaling is Pycsar system, which contains pyrimidine cyclase enzymes that specifically synthesize cCMP and cUMP as second messengers to defense against virus [99]. Some system subtypes in Thoeris and CRISPR-Cas systems encode cyclase domains sometime, which produce diverse cyclic nucleotide and activate downstream of heterotrimeric G-proteins or other receptors [141,142]. Type III CRISPR-Cas systems are among the most widespread and diverse CRISPR-Cas systems [29], where CRISPR effectors specifically recognize and cleave complementary RNA, and further activate Cas10 protein and ancillary proteins which produce cyclic oligoadenylates (cOAs) to activate downstream effectors [142,143].

#### 2.3.4 Other eukaryotic immunity related systems

In addition to the antiviral signaling systems, there are other defense systems similar with eukaryotic immunity, suggesting that these proteins involved in eukaryotic immunity may originate from prokaryotes. For example, Viperin is an antiviral protein stimulated by interferon that effectively inhibits viral release in mammalian cells [144]. Viperin catalyzes the conversion of CTP to ddhCTP, which lacks a hydroxyl group on the 3’ position of the carbon, thus terminates the viral RNA transcription [145]. Recent studies showed that prokaryotic viperins (pVips) can also produce modified ribonucleotides including ddhCTP, ddhGTP, and ddhUTP, providing defense against bacteriophages [145]. Asgard archaea, which is the prokaryote most closely related with eukaryote [146,147], was also found to encode active pVips, suggesting their presence in the Last Eukaryotic Common Ancestor (LECA) [148]. Similar with pVips that deplete the nucleotides essential for replication or transcription, dCTPdeaminase and dGTPase system converts dCTP into deoxy- uracil nucleotides or degrades dGTP into phosphate-free deoxyguanosine, respectively [124].

The single-gene SEFIR system has likewise been mentioned to mediate the TIR signaling pathway, which protein structure similar with TIR domain [65]. Double-gene Eleos system was thought to be related with dynamins, where LeoBC was found homology to GTPases immunity-associated proteins (GIMAPs) [65,149]. Four-gene ISG15-like system comprises a gene with homology to ISG15 (Interferon-stimulated gene 15), two genes with homology to ubiquitin-conjugating enzymes E1 and E2, and a gene homology to ubiquitin-removing enzymes of the JAB/JAMM family [65,150,151].

### 2.4 Other diverse systems

In addition to the systems mentioned above that can be clearly classified into the three major groups, some of the remaining systems encode other diverse proteins. Four systems are related with proteases, which may degrade proteins involved in cellular metabolism to trigger growth arrest through Abi mechanism. For example, Lit protein in *E. coli* is activated by recognizing the coat protein Gp23 of the T4 phage, and further cleaves the elongation factor EF-Tu to prevents the translation and protein synthesis in host [152]. Borvo system was thought to be functioned by CHAT protease domain to resist T4 phage and some siphoviruses [65]. The Gao_Iet system is composed of two genes, one of which encodes protease, while the other gene is associated with ATPase [62]. Expression of the phage gene was showed toxic in cells containing Borvo or Gao_Iet system, indicating that these systems will kill the infected cells [68]. PD-Lambda-2 system encodes a gene with HigA antitoxin-like domain fused to peptidase domain, which was shown to be toxic [86].

Four systems were related with post-translational modification (PTM) of proteins, including four systems (Stk2, PD-T4-6, Gao_TerY, CapRel) related with kinase. For example, it has been reported that the activation of serine threonine kinase Stk2 will affect the transcription and translation of cells [153], CapRel will pyrophosphorylate the tRNA and further block protein translation [88]. Beside these systems which phosphorylate the subtracts, SoFIC system consists of standalone protein with Fic domain that can carry out PTM through the addition of NMPs, phosphocholine or phosphate to substrate proteins [154].

Lastly, the remainder defense systems include 8 characterized TA systems, and 9 uncharacterized systems. PsyrTA and ShosTA systems both encode phosphoribosyl transferase (PRTase) domain as toxin [37,65]. SanaTA system encodes nucleotidyl transferases (NTPs) domain as toxin [155], which resemble to AbiE TA system [156,157]. The gp29_gp30 system was found in prophage, which can trigger bacterial stress response and abortive infection through (p)ppGpp synthetase domain [158]. The PfiAT system comprises of PfiT toxin and PfiA antitoxin, where PfiT directly binds to PfiA and functions as a corepressor of PfiA for the TA operon [159]. PD-T4-10 [86] and Rst_gop_beta_cll [37] TA systems are encoded by two or three overlapping genes with unknown function, respectively.

Dodola system consists of a DUF6414 protein [65], and another protein related with ClpB, which is a key chaperone participates in bacterial survival under stress [160]. PD-Lambda-6 system is related with lipoprotein [86]. Gao_Ppl system consists of standalone protein with both polymerase/histidinol phosphatase (PHP) and ATPase domains (may be nucleic acids related) [62]. Other seven systems, including AbiC, AbiG, AbiH, AbiN, Aditi, Rst_3HP, and Rst_DUF4238 systems could not be classified into a clearly known domain family so far.

## 3. Distribution of defense systems among bacteria, archaea, and phage

### 3.1 Distribution of defense systems in bacteria

We selected and analyzed 60,791 bacterial genomes from The Genome Taxonomy Database (GTDB) [161] until August 2023 to comprehensively analyze the distribution of defense systems in bacteria. To ensure the reliability of the data, the top 49 abundant bacterial phyla with more than 40 genomes were selected for analysis. Overall, 330,743 defense systems were detected within these genomes, with an average of 5.44 defense systems per bacterial genome. Due to some of these genomes are obtained from metagenomics and could be incomplete, we further calculated the occurrence frequency of defense systems by the number of defense system per million base pairs (Mbp), instead of per genome. In the heat map, the systems are arranged according to the previous classification above. Systems with the same structural domains or functions are placed nearby.

Almost all defense systems (except Rst_2TM_1TM_TIR, Rst_Old_Tin) can be detected in our bacterial genomes, with RM, Cas, SoFIC, Wadjet, AbiE, Septu, AbiD, and CBASS are most frequently distributed (Figure 2a, Supplementary Figure 1a). Among them, the occurrence frequency of RM and Cas systems are 0.67 and 0.15 per Mbp, respectively. The remaining 6 most frequently distributed systems are between 0.05-0.03 per Mbp. For the diversity of defense systems, in the phyla of Dependentiae, Aquificota, and Bipolaricaulota, the diversity of systems is lowest, with less than 20 types of systems be detected. While in phyla Proteobacteria, Bacteroidota, and Firmicutes, they encode most diverse systems, with more than 110 different types (Supplementary Figure 2). Based on the previous categorization of the systems, we observed that the system consisting of the nuclease or helicase domains has a large distribution. The systems which directly block the DNA replication, RNA transcription or translation distribute rare (color yellow in Figure 1). Also, some systems related with membrane (color blue in Figure 1) and the systems boxed in “Other” (color gray in Figure 1) are thinly distributed.

**Figure 2.**
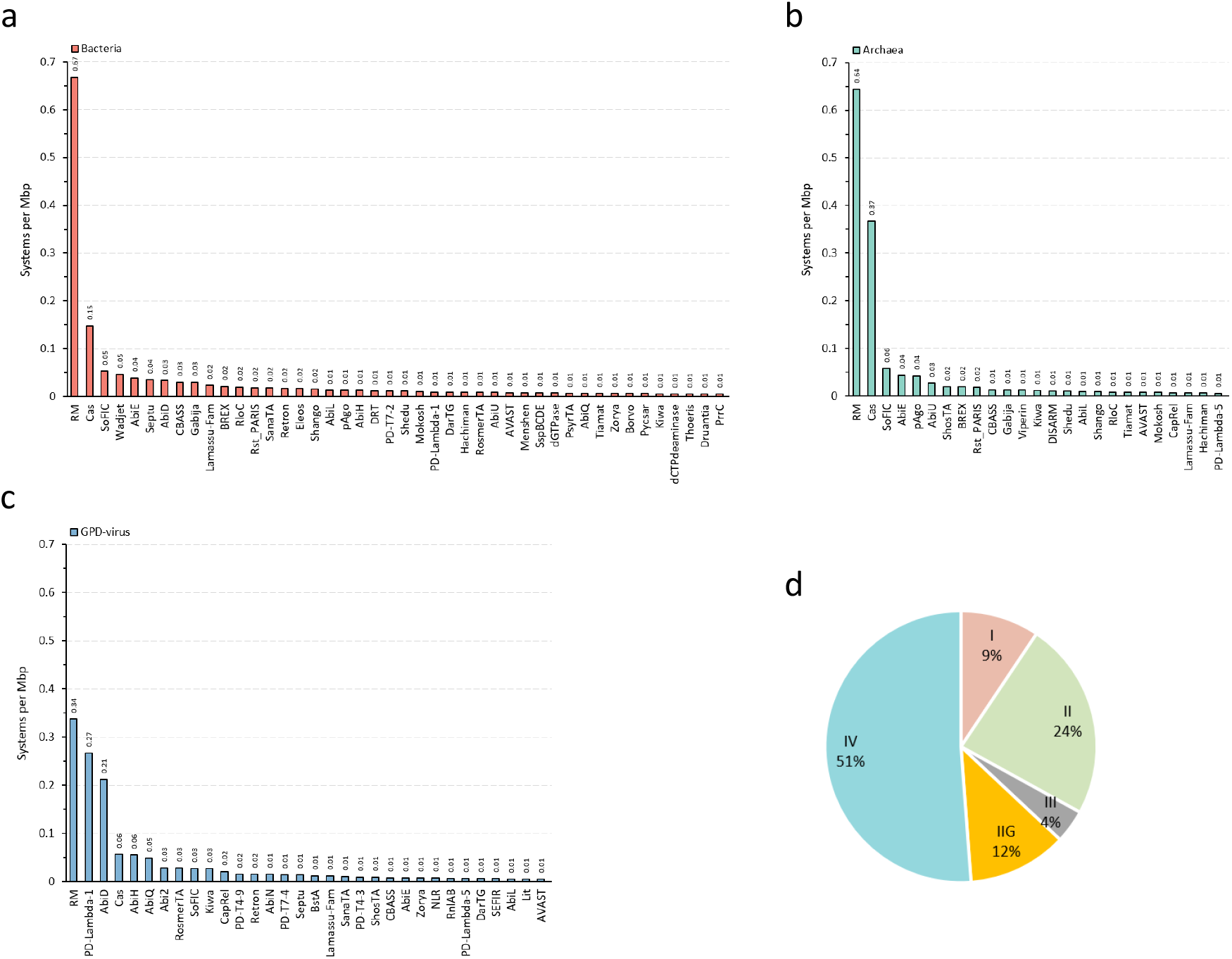
The occurrence frequency of defense systems among bacteria, archaea, and virus domains. **a**, Number of defense systems per Mbp in bacterial genomes. **b**, Number of defense systems per Mbp in archaeal genomes. **c**, Number of defense systems per Mbp in viral genomes. The defense systems with occurrence frequency larger than 0.05 are shown. Defense systems in the x-axis are listed in descending order of number. Frequency is retained to two decimal places. **d**, Distribution of different subtypes of RM system in viral genomes. N=927.

The analysis show that some defense systems are less distributed across different bacterial phyla. Five systems (Pif, PD-Lambda-3, PD-Lambda-4, Rst_gop_beta_cll, Rst_RT-nitrilase-Tm) distribute only in phylum Proteobacteria. AbiB system only distributes in phylum Firmicutes. AbiK, gp29_gp30, MqsRAC, AbiI, Mok_Hok_Sok, and RADAR systems are also less distributed within less than 5 bacterial phyla. These systems with low distribution across bacterial domain also show lower occurrence frequency than 0.0002 per Mbp. Notably, RM, Cas, and SoFIC systems distributed across all bacterial phyla, with occurrence frequency higher than 0.05 per Mbp.

Given that the number of genomes of some bacterial phyla are below 100, some defense systems may encode within bacterial phyla that have not yet been included. From the distribution pattern of defense systems, it can be found that the diversity of defense systems is somewhat positively correlates with the number of genomes within each phylum, suggesting that the analysis of defense systems may be affected by the number of genomes.

### 3.2 Distribution of defense systems in archaea

Among the genomes of more than 60,000 prokaryotes in the GTDB database, 3,412 genomes are archaeal genomes. We first analyzed theses genome as a representative. Overall, 8,403 defense systems were detected in these genomes, with an average of 2.46 defense systems per genome. However, to improve the reliability of the data, we again downloaded over 12,000 archaeal genomes from National Center for Biotechnology Information (NCBI) database, and further assigned their taxonomy through GTDB standard. We removed about 3,000 genomes with unclear taxonomy or with low integrity, finally retained 8,973 archaeal genomes for further analysis. According to the previous classified standard [162], these archaeal genomes were divided into four superfamilies, including Asgard, TACK, DPANN, and Euryarchaeota groups. We also used new GTDB classification standard to divide them into 45 phyla, classes, or orders for comparison.

A total number of 29,028 defense systems (100 types) are detected, with an average of 3.24 defense systems per genome. In general, the defense system of archaeal domain showed lower diversity and abundance than bacterial domain. For the diversity, all 45 archaeal taxa encode RM system, while 42 and 39 taxa encode Cas and pAgo systems respectively (Supplementary Figure 3). AbiE, Shedu, SoFIC, ShosTA, and Gabija systems are also widely distributed in more than 25 archaeal taxa. However, 16 systems (dGTPase, ISG15-like, Bunzi, etc.) only distributed in one archaeal taxon, while 44 systems distributed in archaeal taxa less than 10, including Thoeris, Gasdermin, and NLR systems which are related with eukaryotic immunity. The distribution pattern based on the previous categorization of the systems are similar to that of bacteria, where the system consisting of the nuclease or helicase domains has a large distribution.

Based on the classification of the archaeal superphylum groups, Euryarchaeota contained 91 types of different systems, while 69, 61, 60 types in DPANN, TACK, and Asgard group, respectively (Figure 3). Among them, 11 systems (Dodola, Aditi, Uzume, etc.) distribute only in Euryarchaeota group, while 3 (Dazbog, PD-T7-1, Gao_Qat), 2 (PD-T7-3, PD-Lambda-2), and 1 (dGTPase) system only in DPANN, TACK, and Asgard groups, respectively. Thirty systems (RADAR, NixI, BstA, etc.) were not detected in archaea, indicating that these 30 systems may specific in bacteria.

**Figure 3.**
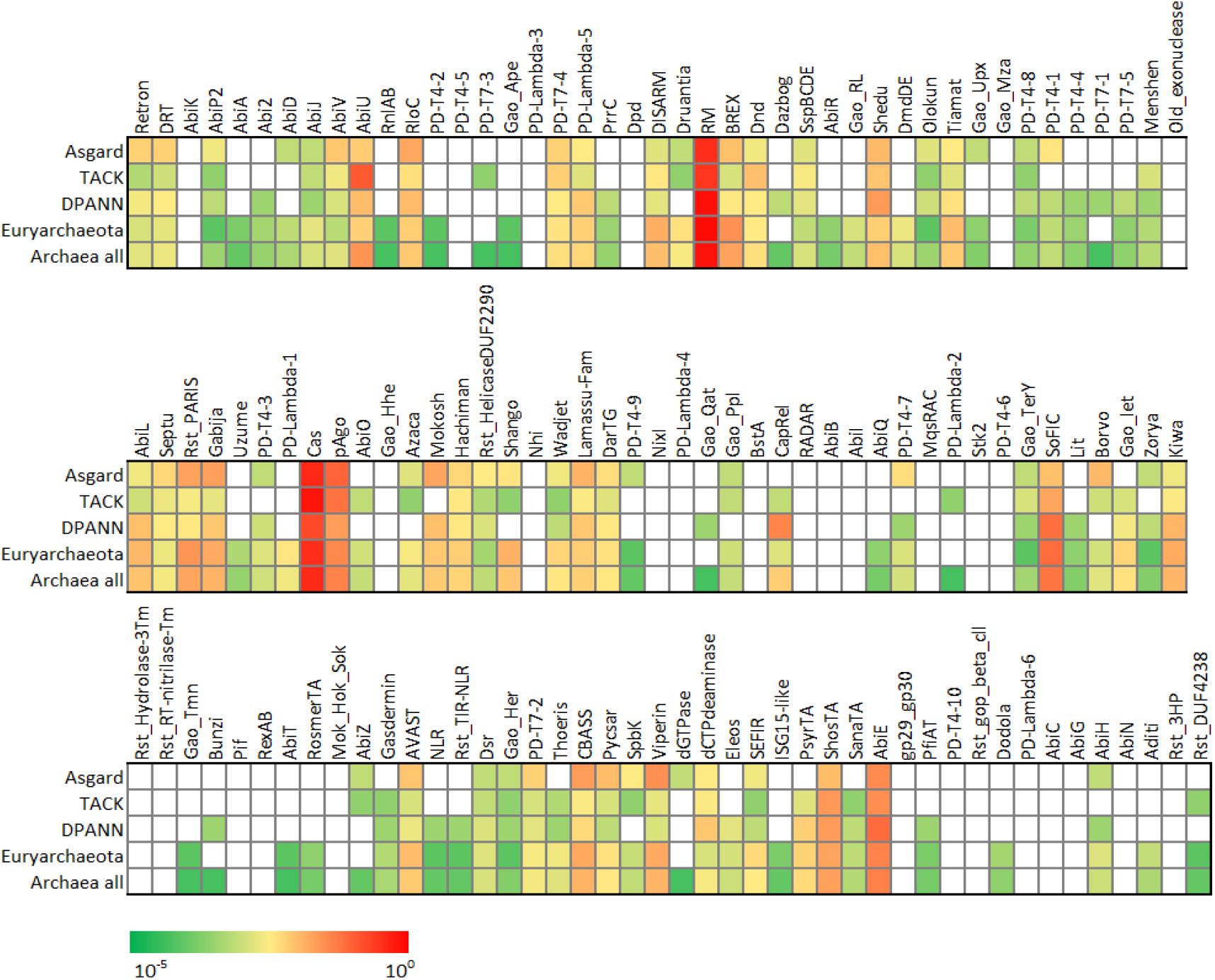
Distribution of defense systems among different archaeal superfamilies. Data are shown by the number of systems per Mbp, and are demonstrated in logarithmic scale colormap.

For the occurrence frequency, we found that RM, Cas, SoFIC, AbiE, pAgo, AbiU, ShosTA, and BREX systems are the 8 most frequently occur systems, with frequency from 0.64-0.02 per Mbp (Figure 2b, Supplementary Figure 1b). The average number of defense systems per Mbp in the Asgard superphylum is 1.14, while in the TACK group it is 1.22 and in the DPANN group it is 1.28. The frequency of these systems in the Euryarchaeota is the highest, reaching 1.59 per Mbp. Notably, we also found some systems (Viperin, CBASS, pAgo, etc.) related with eukaryotic immunity are distribute in Asgard group, which might provide some hints to the eukaryogenesis as Asgard archaea is believed to be the prokaryotes most closely related with eukaryotes so far [146,147].

### 3.3 Distribution of defense systems in phage

It has been reported that some prophages encoded in bacterial genomes carried defense systems to prevent the infection from other phages. Therefore, we analyzed the distribution of defense systems in phage genomes. We first analyzed 11,350 complete genomes of isolated phages from NCBI database (up to August 2023). The results showed that 380 defense systems existed in these genomes, with frequency of 0.033 system per genome, including 44 different types of systems (results not shown). Among them, AbiD, Cas, and RM system were dominant. Given that the genome size of phages is relatively small, generally between 10-100 kb, to reduce the error caused by the small amount of data, we further selected 142,835 high quality phage genomes from Gut Phage Database (GPD) [163].

The results showed that 7,609 defense systems are encoded in GPD genomes, with an average of 0.053 systems per genome. Totally 103 types of defense systems encoded in the genome of these enteroviruses, suggesting that most of the defense systems currently found are encoded by phages too. The occurrence frequency of RM, PD Lambda-1, and AbiD systems are the highest among these phages, reaching 0.34, 0.27, and 0.21 per Mbp, respectively (Figure 2c, Supplementary Figure 1c). Other 7 enriched defense systems are Cas, AbiH, AbiQ, Abi2, RosmerTA, SoFIC, and Kiwa systems, with frequency between 0.06 to 0.03 per Mbp. These defense systems encoded by phages are mainly single-gene system in Abi systems, but there are also some multiple-genes systems, such as Cas, CBASS, Lamassu-Fam, etc.

We further analyzed the most enriched RM systems encoded in phage genomes, which are also the most abundant systems in bacteria and archaea. The results showed that the proportion of type I RM system, which forms multi-subunit complex, accounts for 9% in gut phage genomes (Figure 2d). While type II RM system accounting 24%, which is thought to be the most common system in prokaryotes that consists of separate REase and MTase. Type IIG is a subtype of the type II system, which have REase and MTase domains within a single protein rather than being separated. The proportion of this system accounts for 12% in phage genomes. The type III RM system accounts for 4% in phage genomes. Notably, the type IV RM system accounts for 51% in phage genomes, which has only one REase without a MTase. Consequently, the type IV RM system is dominant among gut phage genomes. This type of RM system has a smaller size compared to others and is suitable for existence in phages with hypercompact genomes.

To better understand the distribution of defense systems between different phages and their hosts. We further classified these phages into known viral taxonomy according to the International Committee on Taxonomy of Viruses (ICTV), as well as assigned their hosts. For the 11.6% of phages (16,636) that could be classified into known viral taxonomy, we detected defense systems mainly in Podoviridae, Siphovirida, and Myoviridae morphotype families (Figure 4a). Myoviridae harbor a significantly higher defense systems frequency, with an average of 2.5 systems per Mbp, while 1.6 of Siphovirida, and 0.4 of Podoviridae. At the same time, Myoviridae also encode more diverse defense systems (58 types) than Siphovirida (37 types) and Myoviridae (17 types). Among them, Retron, Septu, Rst_3HP, AbiU and PD-T4-3 are specific in Myoviridae with higher prevalence.

**Figure 4.**
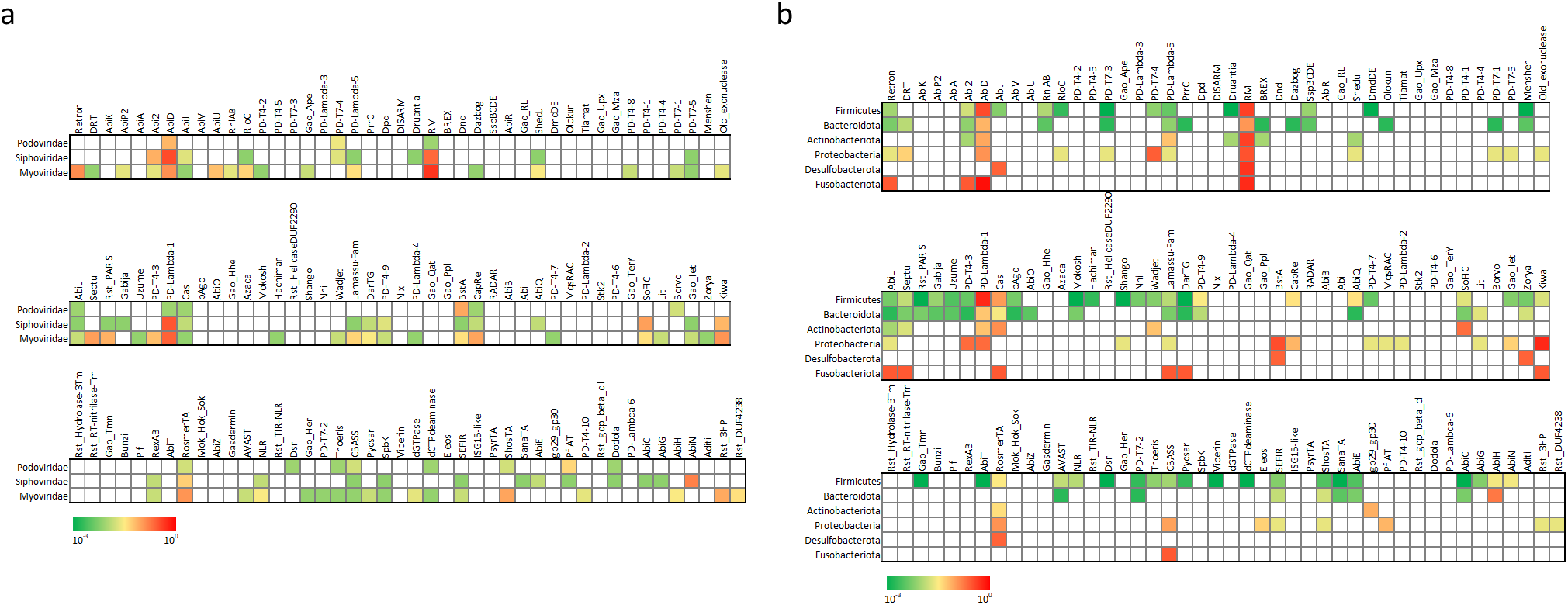
Insight into the distribution of defense systems in different phage taxonomy and hosts. **a**, Distribution of defense systems in Podoviridae, Siphovirida, and Myoviridae families. **b**, Distribution of defense systems encoded by phages with different bacterial hosts. Hosts are distributed in phylum Firmicutes, Bacteroidota, Actinobacteriota, Proteobacteria, Desulfobacterota, or Fusobacteriota. Data are shown by the number of systems per Mbp and demonstrated in logarithmic scale colormap.

For the 28.6% of phages (40,919) which had predicted bacterial hosts, we detected significant higher frequency of defense systems of phages that infected Fusobacteriota, with 3.8 systems per Mbp in comparison with less than 3 for the other phyla. The phages that infected Bacteroidota harbor less defense systems, with an average of 0.6 systems per Mbp. Although Fusobacteriota harbor higher frequency, only 11 kinds systems were encoded with a low diversity. Meanwhile, Bacteroidota harbor lower defense systems rate, but it has higher diversity with 41 different defense systems. The diversity of defense systems in phages hosted by Desulfobacterota is relatively low, only five types of systems (AbiJ, BstA, RM, RosmerTA, Zorya) have been detected. Some defense systems only distribute within specific host phages. For example, AbiT, DmdDE, dCTPdeaminase, Gao_Tmn, Hachiman, Nhi, and NLR systems only present in phages hosted by Firmicutes (Figure 4b). Due to the difficulty in predicting the host or classification of most phages, the distribution of some defense systems is still unpredictable. Future research on validating these phage-encoded defense systems will be interesting.

### 3.4 Comparison between bacteria, archaea, and phage

To find the difference between the distribution of defense systems among bacteria, archaea, and phage, we first compared their occurrence frequency of total defense systems. The results show that an average of 1.59 systems per Mbp in bacterial genomes, 1.45 systems per Mbp in archaeal genomes, while 1.41 systems per Mbp in viral genomes. We noticed that RM, Cas, RM, and SoFIC systems are all relatively enriched in bacteria, archaea, and phage (Figure 5a, b).

**Figure 5.**
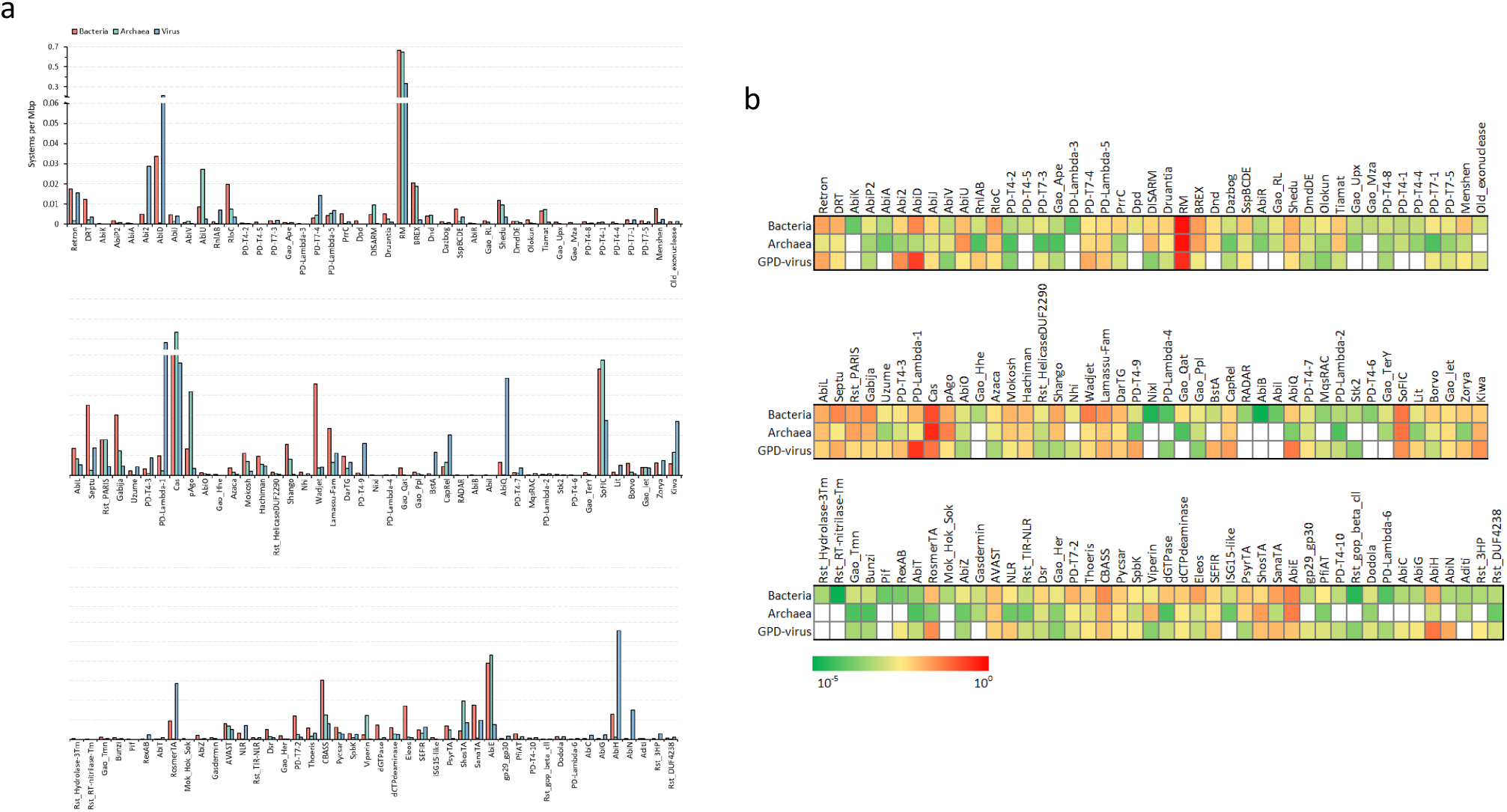
Comparison of defense systems encoded by bacteria, archaea and virus. Data are shown by the number of systems per Mbp, and are demonstrated in **a**, bar chat, and **b**, logarithmic scale colormap. The y-axis of bar chat is cut from 0.06 to 0.2 for data visualization.

We further focused on the systems with significant differences among these three domains. The occurrence frequency of 19 systems of archaea is higher than that of bacteria, including Kiwa, DISARM, AbiV, etc. Specially, the frequency of archaeal AbiU, Cas, pAgo, ShosTA, and Viperin systems are significantly higher than bacteria, indicating that some of these defense systems may initially originate from archaea. The occurrence frequency of 39 systems of phages is higher than that of bacteria, including CapRel, AbiH, PD- T7-4, etc. Specially, the frequency of PD-Lambda-1, PD-T4-9, RnlAB, Abi2, AbiD, and AbiQ are significantly higher than both bacteria and archaea. Given that these systems are mostly composed of single-gene and do not overly burden the viral genome, they may be preserved in phages and enhance virus exclusivity. Overall, most defense systems that have been discovered so far, the abundance of bacteria is generally higher than that of archaea and phage. This may be due to the bias introduced by the use of bacteria as model strains for most studies. Our results also indicated that numerous phages encode their own defense systems that may prevent superinfection by other phages and maintain host fitness.

## 4 Conclusion

In this paper, we categorized the defense systems by their structural functional domains. We propose that diverged defense systems are originated from simple function domains with sensors and effectors, which gives us an idea of finding novel defense systems. They combined and evolved to different systems in different species through further horizontal gene transfer. These functional domains protect the prokaryotes from phage infection. The assembly of different components results in different sensing and effect modes, and different defense systems respond differently to MGEs. Also, through bioinformatics analysis and previous studies, we found that eukaryotic and prokaryotic immunity are significantly related to apoptosis, signal transduction, and homology of chromosomal structure stabilizing proteins.

This study analyzed the distribution of defenses systems with 212,599 genomes, which should be the largest analyzed genomes to date. The results show that archaeal and viral genomes harbor high frequency of defense systems, which has not reported in the previous study. Since most of the previous studies used bacteria as model organisms, this may have resulted in some systems specific to archaea not being identified. We also suspect that phages may acquire some defense genes from the host, using the same systems to resist invasion by other phages. It may through phage infection that the defense systems can spread among different species.

## Author contribution

Jinquan Li and Meng Li drafted the manuscript and figures. Jinquan Li, Jiazheng Gu, and Runyue Xia performed data analyses and wrote the manuscript. Meng Li supervised and funded this project. All authors have read the final manuscript and approved it for publication.

## Acknowledgements

We are grateful to Professor Peter Fineran in University of Otago for critical reading and giving useful suggestion of the manuscript. This work was supported by the National Key Research and Development Program of China (2022YFA0912200), the National Natural Science Foundation of China (32225003, 32393970, 32393971), the Shenzhen Medical Research Fund (B2301005), Guangdong Major Project of Basic and Applied Basic Research (2023B0303000017), Shenzhen University 2035 Program for Excellent Research (2022B002) and the Synthetic Biology Research Center of Shenzhen University.

## Data availability statement

Supplementary information is available online DOI.

## Reference

1. Makarova KS, Anantharaman V, Aravind L, Koonin E V.: Live virus-free or die: coupling of antivirus immunity and programmed suicide or dormancy in prokaryotes. Biol Direct 2012, 7:1.

2. Touchon M, Bernheim A, Rocha EPC: Genetic and life-history traits associated with the distribution of prophages in bacteria. ISME J 2016, doi:10.1038/ismej.2016.47.

3. Bernheim A, Sorek R: The pan-immune system of bacteria: antiviral defence as a community resource. Nat Rev Microbiol 2020, 18:113–119.

4. Pfeifer E, Bonnin RA, Rocha EPC: Phage-Plasmids Spread Antibiotic Resistance Genes through Infection and Lysogenic Conversion. MBio 2022, doi:10.1128/mbio.01851-22.

5. Brown-Jaque M, Calero-Cáceres W, Muniesa M: Transfer of antibiotic-resistance genes via phage-related mobile elements. Plasmid 2015, doi:10.1016/j.plasmid.2015.01.001.

6. van Houte S, Buckling A, Westra ER: Evolutionary Ecology of Prokaryotic Immune Mechanisms. Microbiol Mol Biol Rev 2016, 80:745–763.

7. Hampton HG, Watson BNJ, Fineran PC: The arms race between bacteria and their phage foes. Nature 2020, 577:327–336.

8. Tock MR, Dryden DTF: The biology of restriction and anti-restriction. Curr Opin Microbiol 2005, 8:466–472.

9. Brouns SJJ, Jore MM, Lundgren M, Westra ER, Slijkhuis RJH, Snijders APL, Dickman MJ, Makarova KS, Koonin E V., Van Der Oost J: Small CRISPR RNAs guide antiviral defense in prokaryotes. Science (80- ) 2008, 321:960–964.

10. Roberts RJ, Vincze T, Posfai J, Macelis D: REBASE-a database for DNA restriction and modification: Enzymes, genes and genomes. Nucleic Acids Res 2015, 43:D298–D299.

11. Koonin E V, Makarova KS, Wolf YI: Evolutionary Genomics of Defense Systems in Archaea and Bacteria. Annu Rev Microbiol 2017, 71:233–261.

12. Lopatina A, Tal N, Sorek R: Abortive Infection: Bacterial Suicide as an Antiviral Immune Strategy. Annu Rev Virol 2020, 7:371–384.

13. Leroux M, Laub MT: Toxin-Antitoxin Systems as Phage Defense Elements. Annu Rev Microbiol 2022, doi:10.1146/annurev-micro-020722-013730.

14. Makarova KS, Wolf YI, Snir S, Koonin E V.: Defense Islands in Bacterial and Archaeal Genomes and Prediction of Novel Defense Systems. J Bacteriol 2011, 193:6039–6056.

15. Makarova KS, Wolf YI, Koonin E V.: Comparative genomics of defense systems in archaea and bacteria. Nucleic Acids Res 2013, 41:4360–4377.

16. Georjon H, Bernheim A: The highly diverse antiphage defence systems of bacteria. Nat Rev Microbiol 2023, 21:686–700.

17. Tesson F, Hervé A, Mordret E, Touchon M, D’Humières C, Cury J, Bernheim A: Systematic and quantitative view of the antiviral arsenal of prokaryotes. Nat Commun 2022, 13:2561.

18. Aravind L, Koonin E V.: DNA polymerase β-like nucleotidyltransferase superfamily: Identification of three new families, classification and evolutionary history. Nucleic Acids Res 1999, doi:10.1093/nar/27.7.1609.

19. Anantharaman V, Makarova KS, Burroughs AM, Koonin E V., Aravind L: Comprehensive analysis of the HEPN superfamily: Identification of novel roles in intra-genomic conflicts, defense, pathogenesis and RNA processing. Biol Direct 2013, 8:1.

20. Pillon MC, Stanley RE: Nuclease integrated kinase super assemblies (NiKs) and their role in RNA processing. Curr Genet 2018, doi:10.1007/s00294-017-0749-9.

21. Ricagno S, Egloff MP, Ulferts R, Coutard B, Nurizzo D, Campanacci V, Cambillau C, Ziebuhr J, Canard B: Crystal structure and mechanistic determinants of SARS coronavirus nonstructural protein 15 define an endoribonuclease family. Proc Natl Acad Sci U S A 2006, doi:10.1073/pnas.0601708103.

22. Coelho DS, Domingos PM: Physiological roles of regulated Ire1 dependent decay. Front Genet 2014, doi:10.3389/fgene.2014.00076.

23. Pillon MC, Sobhany M, Borgnia MJ, Williams JG, Stanley RE, Baker D: Grc3 programs the essential endoribonuclease Las1 for specific RNA cleavage. Proc Natl Acad Sci U S A 2017, doi:10.1073/pnas.1703133114.

24. Davidov E, Kaufmann G: RloC: A wobble nucleotide-excising and zinc-responsive bacterial tRNase. Mol Microbiol 2008, doi:10.1111/j.1365-2958.2008.06387.x.

25. Meineke B, Shuman S: Structure-function relations in the NTPase domain of the antiviral tRNA ribotoxin Escherichia coli PrrC. Virology 2012, doi:10.1016/j.virol.2012.02.008.

26. Yao J, Zhen X, Tang K, Liu T, Xu X, Chen Z, Guo Y, Liu X, Wood TK, Ouyang S, et al.: Novel polyadenylylation-dependent neutralization mechanism of the HEPN/MNT toxin/antitoxin system. Nucleic Acids Res 2020, 48:11054–11067.

27. Pillon MC, Gordon J, Frazier MN, Stanley RE: HEPN RNases–an emerging class of functionally distinct RNA processing and degradation enzymes. Crit Rev Biochem Mol Biol 2021, doi:10.1080/10409238.2020.1856769.

28. Koga M, Otsuka Y, Lemire S, Yonesaki T: Escherichia coli rnlA and rnlB compose a novel toxin-antitoxin system. Genetics 2011, doi:10.1534/genetics.110.121798.

29. Makarova KS, Wolf YI, Iranzo J, Shmakov SA, Alkhnbashi OS, Brouns SJJ, Charpentier E, Cheng D, Haft DH, Horvath P, et al.: Evolutionary classification of CRISPR–Cas systems: a burst of class 2 and derived variants. Nat Rev Microbiol 2020, 18:67–83.

30. Sironi G: Mutants of Escherichia coli unable to be lysogenized by the temperate bacteriophage P2. Virology 1969, doi:10.1016/0042-6822(69)90196-2.

31. Schiltz CJ, Lee A, Partlow EA, Hosford CJ, Chappie JS: Structural characterization of Class 2 OLD family nucleases supports a two-metal catalysis mechanism for cleavage. Nucleic Acids Res 2019, 47:9448–9463.

32. Schiltz CJ, Adams MC, Chappie JS: The full-length structure of Thermus scotoductus OLD defines the ATP hydrolysis properties and catalytic mechanism of Class 1 OLD family nucleases. Nucleic Acids Res 2020, 48:2762–2776.

33. Doron S, Melamed S, Ofir G, Leavitt A, Lopatina A, Keren M, Amitai G, Sorek R: Systematic discovery of antiphage defense systems in the microbial pangenome. Science (80- ) 2018, 359:0–12.

34. Antine SP, Johnson AG, Mooney SE, Leavitt A, Mayer ML, Yirmiya E, Amitai G, Sorek R, Kranzusch PJ: Structural basis of Gabija anti-phage defence and viral immune evasion. Nature 2024, 625:360–365.

35. Li J, Cheng R, Wang Z, Yuan W, Xiao J, Zhao X, Du X, Xia S, Wang L, Zhu B, et al.: Structures and activation mechanism of the Gabija anti-phage system. Nature 2024, 629.

36. Cheng R, Huang F, Lu X, Yan Y, Yu B, Wang X, Zhu B: Prokaryotic Gabija complex senses and executes nucleotide depletion and DNA cleavage for antiviral defense. Cell Host Microbe 2023, doi:10.1016/j.chom.2023.06.014.

37. Rousset F, Depardieu F, Miele S, Dowding J, Laval AL, Lieberman E, Garry D, Rocha EPC, Bernheim A, Bikard D: Phages and their satellites encode hotspots of antiviral systems. Cell Host Microbe 2022, 30:740–753.e5.

38. Deep A, Liang Q, Enustun E, Pogliano J, Corbett KD: Architecture and infection-sensing mechanism of the bacterial PARIS defense system. bioRxiv 2024,

39. Atanasiu C, Su TJ, Sturrock SS, Dryden DTF: Interaction of the ocr gene 0.3 protein of bacteriophage T7 with EcoKl restriction/modification enzyme. Nucleic Acids Res 2002, doi:10.1093/nar/gkf518.

40. Isaev A, Drobiazko A, Sierro N, Gordeeva J, Yosef I, Qimron U, Ivanov N V., Severinov K: Phage T7 DNA mimic protein Ocr is a potent inhibitor of BREX defence. Nucleic Acids Res 2020, doi:10.1093/NAR/GKAA290.

41. Payne LJ, Todeschini TC, Wu Y, Perry BJ, Ronson CW, Fineran PC, Nobrega FL, Jackson SA: Identification and classification of antiviral defence systems in bacteria and archaea with PADLOC reveals new system types. Nucleic Acids Res 2021, 49:10868–10878.

42. Millman A, Bernheim A, Stokar-Avihail A, Fedorenko T, Voichek M, Leavitt A, Sorek R: Bacterial retrons function in anti-phage defense. Cell 2020, doi:10.1101/2020.06.21.156273.

43. Mestre MR, González-Delgado A, Gutiérrez-Rus LI, Martínez-Abarca F, Toro N: Systematic prediction of genes functionally associated with bacterial retrons and classification of the encoded tripartite systems. Nucleic Acids Res 2020, doi:10.1093/nar/gkaa1149.

44. Li, Shen Z, Zhang M, Yang XY, Cleary SP, Xie J, Marathe IA, Kostelic M, Greenwald J, Rish AD, et al.: PtuA and PtuB assemble into an inflammasome-like oligomer for anti-phage defense. Nat Struct Mol Biol 2024, 31:413–423.

45. Aggarwal AK: Structure and function of restriction endonucleases. Curr Opin Struct Biol 1995, doi:10.1016/0959-440X(95)80004-K.

46. Kovall RA, Matthews BW: Structural, functional, and evolutionary relationships between λ- exonuclease and the type II restriction endonucleases. Proc Natl Acad Sci U S A 1998, doi:10.1073/pnas.95.14.7893.

47. Hickman AB, Li Y, Mathew SV, May EW, Craig NL, Dyda F: Unexpected structural diversity in DNA recombination: The restriction endonuclease connection. Mol Cell 2000, doi:10.1016/S1097-2765(00)80267-1.

48. Nishino T, Komori K, Ishino Y, Morikawa K: X-ray and biochemical anatomy of an archaeal XPF/Rad1/Mus81 family nuclease: Similarity between its endonuclease domain and restriction enzymes. Structure 2003, doi:10.1016/S0969-2126(03)00046-7.

49. Belfort M, Weiner A: Another bridge between kingdoms: TRNA splicing in Archaea and Eukaryotes. Cell 1997, doi:10.1016/S0092-8674(00)80287-1.

50. Laganeckas M, Margelevičius M, Venclovas Č: Identification of new homologs of PD-(D/E)XK nucleases by support vector machines trained on data derived from profile–profile alignments. Nucleic Acids Res 2011, 39:1187–1196.

51. Kosinski J, Feder M, Bujnicki JM: The PD-(D/E)XK superfamily revisited: Identification of new members among proteins involved in DNA metabolism and functional predictions for domains of (hitherto) unknown function. BMC Bioinformatics 2005, doi:10.1186/1471-2105-6-172.

52. Steczkiewicz K, Muszewska A, Knizewski L, Rychlewski L, Ginalski K: Sequence, structure and functional diversity of PD-(D/E)XK phosphodiesterase superfamily. Nucleic Acids Res 2012, 40:7016–7045.

53. Kowalski JC, Belfort M, Stapleton MA, Holpert M, Dansereau JT, Pietrokovski S, Baxter SM, Derbyshire V: Configuration of the catalytic GIY-YIG domain of intron endonuclease I-Tevl: Coincidence of computational and molecular findings. Nucleic Acids Res 1999, doi:10.1093/nar/27.10.2115.

54. Aravind L, Walker DR, Koonin E V.: Conserved domains in DNA repair proteins and evolution of repair systems. Nucleic Acids Res 1999, doi:10.1093/nar/27.5.1223.

55. Bujnicki JM, Rychlewski L, Radlinska M: Polyphyletic evolution of type II restriction enzymes revisited: Two independent sources of second-hand folds revealed. Trends Biochem Sci 2001, doi:10.1016/S0968-0004(00)01690-X.

56. Kleinstiver BP, Wolfs JM, Edgell DR: The monomeric GIY-YIG homing endonuclease I-BmoI uses a molecular anchor and a flexible tether to sequentially nick DNA. Nucleic Acids Res 2013, doi:10.1093/nar/gkt186.

57. Dunin-Horkawicz S, Feder M, Bujnicki JM: Phylogenomic analysis of the GIY-YIG nuclease superfamily. BMC Genomics 2006, doi:10.1186/1471-2164-7-98.

58. González-Delgado A, Mestre MR, Martínez-Abarca F, Toro N: Prokaryotic reverse transcriptases: From retroelements to specialized defense systems. FEMS Microbiol Rev 2021, 45:1–19.

59. Fortier LC, Bouchard JD, Moineau S: Expression and site-directed mutagenesis of the lactococcal abortive phage infection protein AbiK. J Bacteriol 2005, doi:10.1128/JB.187.11.3721-3730.2005.

60. Dinsmore PK, Klaenhammer TR: Molecular characterization of a genomic region in a Lactococcus bacteriophage that is involved in its sensitivity to the phage defense mechanism AbiA. J Bacteriol 1997, doi:10.1128/jb.179.9.2949-2957.1997.

61. Odegrip R, Nilsson AS, Haggård-Ljungquist E: Identification of a gene encoding a functional reverse transcriptase within a highly variable locus in the P2-like coliphages. J Bacteriol 2006, 188:1643– 1647.

62. Gao L, Altae-Tran H, Böhning F, Makarova KS, Segel M, Schmid-Burgk JL, Koob J, Wolf YI, Koonin E V., Zhang F: Diverse enzymatic activities mediate antiviral immunity in prokaryotes. Science (80- ) 2020, 369:1077–1084.

63. Singleton MR, Dillingham MS, Wigley DB: Structure and mechanism of helicases and nucleic acid translocases. Annu Rev Biochem 2007, doi:10.1146/annurev.biochem.76.052305.115300.

64. Tuck OT, Adler BA, Armbruster EG, Lahiri A, Hu JJ, Zhou J, Pogliano J, Doudna JA: Hachiman is a genome integrity sensor. bioRxiv 2024, 2024:2024.02.29.582594.

65. Millman A, Melamed S, Leavitt A, Doron S, Bernheim A, Hör J, Garb J, Bechon N, Brandis A, Lopatina A, et al.: An expanded arsenal of immune systems that protect bacteria from phages. Cell Host Microbe 2022, 30:1556–1569.e5.

66. Oliveira PH, Touchon M, Rocha EPC: The interplay of restriction-modification systems with mobile genetic elements and their prokaryotic hosts. Nucleic Acids Res 2014, doi:10.1093/nar/gku734.

67. Dimitriu T, Szczelkun MD, Westra ER: Evolutionary Ecology and Interplay of Prokaryotic Innate and Adaptive Immune Systems. Curr Biol 2020, 30:R1189–R1202.

68. Stokar-Avihail A, Fedorenko T, Hör J, Garb J, Leavitt A, Millman A, Shulman G, Wojtania N, Melamed S, Amitai G, et al.: Discovery of phage determinants that confer sensitivity to bacterial immune systems. Cell 2023, doi:10.1016/j.cell.2023.02.029.

69. Goldfarb T, Sberro H, Weinstock E, Cohen O, Doron S, Charpak-Amikam Y, Afik S, Ofir G, Sorek R: BREX is a novel phage resistance system widespread in microbial genomes. EMBO J 2015, 34:169– 183.

70. Picton DM, Luyten YA, Morgan RD, Nelson A, Smith DL, Dryden DTF, Hinton JCD, Blower TR: The phage defence island of a multidrug resistant plasmid uses both BREX and type IV restriction for complementary protection from viruses. Nucleic Acids Res 2021, doi:10.1093/nar/gkab906.

71. Drobiazko A, Adams M, Skutel M, Kristina P, Matlashov M: Molecular basis of foreign DNA recognition by BREX anti-phage immunity system. *bioRxiv* 2024,

72. Sam C. Went, David M. Picton, Richard D. Morgan, Andrew Nelson, David T. F. Dryden, Darren L. Smith, Nicolas Wenner, Jay C. D. Hinton, Tim R. Blower: Structure and rational engineering of the PglX methyltransferase and specificity factor for BREX phage defence. bioRxiv 2024,

73. Xiong L, Liu S, Chen S, Xiao Y, Zhu B, Gao Y, Zhang Y, Chen B, Luo J, Deng Z, et al.: A new type of DNA phosphorothioation-based antiviral system in archaea. Nat Commun 2019, doi:10.1038/s41467-019-09390-9.

74. Jiang S, Chen K, Wang Y, Zhang Y, Tang Y, Huang W, Xiong X, Chen S, Chen C, Wang L: A DNA phosphorothioation-based Dnd defense system provides resistance against various phages and is compatible with the Ssp defense system. MBio 2023, doi:10.1128/mbio.00933-23.

75. Wang S, Wan M, Huang R, Zhang Y, Xie Y, Wei Y, Ahmad M, Wu D, Hong Y, Deng Z, et al.: SSPABCD-SSPFGH constitutes a new type of DNA phosphorothioate-based bacterial defense system. MBio 2021, doi:10.1128/mBio.00613-21.

76. Xiong X, Wu G, Wei Y, Liu L, Zhang Y, Su R, Jiang X, Li M, Gao H, Tian X, et al.: SspABCD–SspE is a phosphorothioation-sensing bacterial defence system with broad anti-phage activities. Nat Microbiol 2020, doi:10.1038/s41564-020-0700-6.

77. Ofir G, Melamed S, Sberro H, Mukamel Z, Silverman S, Yaakov G, Doron S, Sorek R: DISARM is a widespread bacterial defence system with broad anti-phage activities. Nat Microbiol 2018, 3:90–98.

78. Bravo JPK, Aparicio-Maldonado C, Nobrega FL, Brouns SJJ, Taylor DW: Structural basis for broad anti-phage immunity by DISARM. Nat Commun 2022, doi:10.1038/s41467-022-30673-1.

79. Thiaville JJ, Kellner SM, Yuan Y, Hutinet G, Thiaville PC, Jumpathong W, Mohapatra S, Brochier-Armanet C, Letarov A V., Hillebrand R, et al.: Novel genomic island modifies DNA with 7- deazaguanine derivatives. Proc Natl Acad Sci U S A 2016, doi:10.1073/pnas.1518570113.

80. Meineke B, Shuman S: Determinants of the cytotoxicity of PrrC anticodon nuclease and its amelioration by tRNA repair. RNA 2012, doi:10.1261/rna.030171.111.

81. Uhlmann F: SMC complexes: From DNA to chromosomes. Nat Rev Mol Cell Biol 2016, doi:10.1038/nrm.2016.30.

82. Morozova NE, Potysyeva AS, Vedyaykhv AD: Structural and Functional Features of Bacterial SMC Complexes. Tsitologiya 2023, doi:10.31857/S004137712306007X.

83. Deep A, Gu Y, Gao YQ, Ego KM, Herzik MA, Zhou H, Corbett KD: The SMC-family Wadjet complex protects bacteria from plasmid transformation by recognition and cleavage of closed-circular DNA. Mol Cell 2022, doi:10.1016/j.molcel.2022.09.008.

84. Roisné-Hamelin F, Liu HW, Taschner M, Li Y, Gruber S: Structural basis for plasmid restriction by SMC JET nuclease. Mol Cell 2024, doi:10.1016/j.molcel.2024.01.009.

85. LeRoux M, Srikant S, Teodoro GIC, Zhang T, Littlehale ML, Doron S, Badiee M, Leung AKL, Sorek R, Laub MT: The DarTG toxin-antitoxin system provides phage defence by ADP-ribosylating viral DNA. Nat Microbiol 2022, 7:1028–1040.

86. Vassallo CN, Doering CR, Littlehale ML, Teodoro GIC, Laub MT: A functional selection reveals previously undetected anti-phage defence systems in the E. coli pangenome. Nat Microbiol 2022, doi:10.1038/s41564-022-01219-4.

87. Owen S V., Wenner N, Dulberger CL, Rodwell E V., Bowers-Barnard A, Quinones-Olvera N, Rigden DJ, Rubin EJ, Garner EC, Baym M, et al.: Prophages encode phage-defense systems with cognate self-immunity. Cell Host Microbe 2021, doi:10.1016/j.chom.2021.09.002.

88. Zhang T, Tamman H, Coppieters ’t Wallant K, Kurata T, LeRoux M, Srikant S, Brodiazhenko T, Cepauskas A, Talavera A, Martens C, et al.: Direct activation of a bacterial innate immune system by a viral capsid protein. Nature 2022, doi:10.1038/s41586-022-05444-z.

89. Hurley JM, Woychik NA: Bacterial toxin HigB associates with ribosomes and mediates translation-dependent mRNA cleavage at A-rich sites. J Biol Chem 2009, doi:10.1074/jbc.M109.008763.

90. Yu V, Ronzone E, Lord D, Peti W, Page R: MqsR is a noncanonical microbial RNase toxin that is inhibited by antitoxin MqsA via steric blockage of substrate binding. J Biol Chem 2022, doi:10.1016/j.jbc.2022.102535.

91. Wong F, Amir A: Mechanics and Dynamics of Bacterial Cell Lysis. Biophys J 2019, doi:10.1016/j.bpj.2019.04.040.

92. Zhu X, Shi Z, Mao Y, Lächelt U, Huang R: Cell Membrane Perforation: Patterns, Mechanisms and Functions. Small 2024, doi:10.1002/smll.202310605.

93. Müller DJ, Wu N, Palczewski K: Vertebrate membrane proteins: Structure, function, and insights from biophysical approaches. Pharmacol Rev 2008, doi:10.1124/pr.107.07111.

94. Attwood MM, Schiöth HB: Characterization of Five Transmembrane Proteins: With Focus on the Tweety, Sideroflexin, and YIP1 Domain Families. Front Cell Dev Biol 2021, doi:10.3389/fcell.2021.708754.

95. Kozma D, Simon I, Tusnády GE: PDBTM: Protein data bank of transmembrane proteins after 8 years. Nucleic Acids Res 2013, doi:10.1093/nar/gks1169.

96. De Marothy MT, Elofsson A: Marginally hydrophobic transmembrane α-helices shaping membrane protein folding. Protein Sci 2015, doi:10.1002/pro.2698.

97. Spiess M, Junne T, Janoschke M: Membrane Protein Integration and Topogenesis at the ER. Protein J 2019, doi:10.1007/s10930-019-09827-6.

98. Millman A, Melamed S, Amitai G, Sorek R: Diversity and classification of cyclic-oligonucleotide-based anti-phage signalling systems. Nat Microbiol 2020, doi:10.1038/s41564-020-0777-y.

99. Tal N, Morehouse BR, Millman A, Stokar-Avihail A, Avraham C, Fedorenko T, Yirmiya E, Herbst E, Brandis A, Mehlman T, et al.: Cyclic CMP and cyclic UMP mediate bacterial immunity against phages. Cell 2021, 184:5728–5739.e16.

100. Gao LA, Wilkinson ME, Strecker J, Makarova KS, Macrae RK, Koonin E V., Zhang F: Prokaryotic innate immunity through pattern recognition of conserved viral proteins. Science (80- ) 2022, 377:139–148.

101. Todeschini TC, Wu Y, Naji A, Mondi R, Nobrega FL: Kiwa rescues RecBCD for anti-phage activity. bioRxiv 2023,

102. Kayagaki N, Stowe IB, Lee BL, O’Rourke K, Anderson K, Warming S, Cuellar T, Haley B, Roose-Girma M, Phung QT, et al.: Caspase-11 cleaves gasdermin D for non-canonical inflammasome signalling. Nature 2015, doi:10.1038/nature15541.

103. Shi J, Zhao Y, Wang K, Shi X, Wang Y, Huang H, Zhuang Y, Cai T, Wang F, Shao F: Cleavage of GSDMD by inflammatory caspases determines pyroptotic cell death. Nature 2015, doi:10.1038/nature15514.

104. Saeki N, Kuwahara Y, Sasaki H, Satoh H, Shiroishi T: Gasdermin (Gsdm) localizing to mouse chromosome 11 is predominantly expressed in upper gastroiatestinal tract but significantly suppressed in human gastric cancer cells. Mamm Genome 2000, doi:10.1007/s003350010138.

105. Broz P, Pelegrín P, Shao F: The gasdermins, a protein family executing cell death and inflammation. Nat Rev Immunol 2020, doi:10.1038/s41577-019-0228-2.

106. Johnson AG, Wein T, Mayer ML, Duncan-Lowey B, Yirmiya E, Oppenheimer-Shaanan Y, Amitai G, Sorek R, Kranzusch PJ: Bacterial gasdermins reveal an ancient mechanism of cell death. Science (80- ) 2022, 375:221–225.

107. Hu H, Hughes TCD, Popp PF, Roa-Eguiara A, Martin FJO: Structure and mechanism of Zorya anti-phage defense system. bioRxiv 2023,

108. Parma DH, Snyder M, Sobolevski S, Nawroz M, Brody E, Gold L: The Rex system of bacteriophage λ: Tolerance and altruistic cell death. Genes Dev 1992, doi:10.1101/gad.6.3.497.

109. Montgomery MT, Guerrero Bustamante CA, Dedrick RM, Jacobs-Sera D, Hatfull GF: Yet more evidence of collusion: A new viral defense system encoded by gordonia phage carolann. MBio 2019, doi:10.1128/mBio.02417-18.

110. Mageeney CM, Mohammed HT, Dies M, Anbari S, Cudkevich N, Chen Y, Buceta J, Ware VC: Mycobacterium Phage Butters-Encoded Proteins Contribute to Host Defense against Viral Attack . mSystems 2020, doi:10.1128/msystems.00534-20.

111. Durmaz E, Klaenhammer TR: Abortive phage resistance mechanism AbiZ speeds the lysis clock to cause premature lysis of phage-infected Lactococcus lactis. J Bacteriol 2007, 189:1417–1425.

112. Pecota DC, Wood TK: Exclusion of T4 phage by the hok/sok killer locus from plasmid R1. J Bacteriol 1996, doi:10.1128/jb.178.7.2044-2050.1996.

113. Thisted T, Gerdes K: Mechanism of post-segregational killing by the hok/sok system of plasmid R1. J Mol Biol 1992, doi:10.1016/0022-2836(92)90714-u.

114. Schmitt CK, Kemp P, Molineux IJ: Genes 1.2 and 10 of bacteriophages T3 and T7 determine the permeability lesions observed in infected cells of Escherichia coli expressing the F plasmid gene pifA. J Bacteriol 1991, doi:10.1128/jb.173.20.6507-6514.1991.

115. Cheng X, Wang WF, Molineux IJ: F exclusion of bacteriophage T7 occurs at the cell membrane. Virology 2004, doi:10.1016/j.virol.2004.06.001.

116. Anantharaman V, Iyer LM, Aravind L: Ter-dependent stress response systems: Novel pathways related to metal sensing, production of a nucleoside-like metabolite, and DNA-processing. Mol Biosyst 2012, doi:10.1039/c2mb25239b.

117. Ernits K, Saha CK, Brodiazhenko T, Chouhan B, Shenoy A, Buttress JA, Duque-Pedraza JJ, Bojar V, Nakamoto JA, Kurata T, et al.: The structural basis of hyperpromiscuity in a core combinatorial network of type II toxin–antitoxin and related phage defense systems. Proc Natl Acad Sci U S A 2023, doi:10.1073/pnas.2305393120.

118. Fitzgerald KA, Kagan JC: Toll-like Receptors and the Control of Immunity. Cell 2020, doi:10.1016/j.cell.2020.02.041.

119. Burch-Smith TM, Dinesh-Kumar SP: The functions of plant TIR domains. Sci STKE 2007, doi:10.1126/stke.4012007pe46.

120. North BJ, Verdin E: Sirtuins: Sir2-related NAD-dependent protein deacetylases. Genome Biol 2004, doi:10.1186/gb-2004-5-5-224.

121. Yuan H, Marmorstein R: Structural basis for sirtuin activity and inhibition. J Biol Chem 2012, doi:10.1074/jbc.R112.372300.

122. Wan L, Essuman K, Anderson RG, Sasaki Y, Monteiro F, Chung EH, Nishimura EO, DiAntonio A, Milbrandt J, Dangl JL, et al.: TIR domains of plant immune receptors are NAD+-cleaving enzymes that promote cell death. Science *(80- )* 2019, doi:10.1126/science.aax1771.

123. Horsefield S, Burdett H, Zhang X, Manik MK, Shi Y, Chen J, Qi T, Gilley J, Lai JS, Rank MX, et al.: NAD+ cleavage activity by animal and plant TIR domains in cell death pathways. Science *(80- )* 2019, doi:10.1126/science.aax1911.

124. Ofir G, Herbst E, Baroz M, Cohen D, Millman A, Doron S, Tal N, Malheiro DBA, Malitsky S, Amitai G, et al.: Antiviral activity of bacterial TIR domains via immune signalling molecules. Nature 2021, 600:116–120.

125. Wein T, Sorek R: Bacterial origins of human cell-autonomous innate immune mechanisms. Nat Rev Immunol 2022, 22:629–638.

126. Morehouse BR, Govande AA, Millman A, Keszei AFA, Lowey B, Ofir G, Shao S, Sorek R, Kranzusch PJ: STING cyclic dinucleotide sensing originated in bacteria. Nature 2020, 586:429–433.

127. Koopal B, Potocnik A, Mutte SK, Aparicio-Maldonado C, Lindhoud S, Vervoort JJM, Brouns SJJ, Swarts DC: Short prokaryotic Argonaute systems trigger cell death upon detection of invading DNA. Cell 2022, 185:1471–1486.e19.

128. Koopal B, Mutte SK, Swarts DC: A long look at short prokaryotic Argonautes. Trends Cell Biol 2023, 33:605–618.

129. Kibby EM, Conte AN, Burroughs AM, Nagy TA, Vargas JA, Whalen LA, Aravind L, Whiteley AT: Bacterial NLR-related proteins protect against phage. Cell 2023, 186:2410–2424.e18.

130. Leipe DD, Koonin E V., Aravind L: STAND, a class of P-loop NTPases including animal and plant regulators of programmed cell death: Multiple, complex domain architectures, unusual phyletic patterns, and evolution by horizontal gene transfer. J Mol Biol 2004, doi:10.1016/j.jmb.2004.08.023.

131. Koganti R, Patil CD, Shukla D: STAND alert! Prokaryotic immunity for recognition and defense against bacteriophages. Trends Microbiol 2022, 30:1128–1130.

132. Zaccolo M, Zerio A, Lobo MJ: Subcellular organization of the camp signaling pathway. Pharmacol Rev 2021, doi:10.1124/pharmrev.120.000086.

133. Berridge MJ: Calcium microdomains: Organization and function. Cell Calcium 2006, doi:10.1016/j.ceca.2006.09.002.

134. Murad F: Nitric Oxide: The Coming of the Second Messenger. Rambam Maimonides Med J 2011, doi:10.5041/rmmj.10038.

135. Sun L, Wu J, Du F, Chen X, Chen ZJ: Cyclic GMP-AMP synthase is a cytosolic DNA sensor that activates the type I interferon pathway. Science (80- ) 2013, doi:10.1126/science.1232458.

136. Wu J, Sun L, Chen X, Du F, Shi H, Chen C, Chen ZJ: Cyclic GMP-AMP is an endogenous second messenger in innate immune signaling by cytosolic DNA. Science (80- ) 2013, doi:10.1126/science.1229963.

137. Ablasser A, Goldeck M, Cavlar T, Deimling T, Witte G, Röhl I, Hopfner KP, Ludwig J, Hornung V: CGAS produces a 2′-5′-linked cyclic dinucleotide second messenger that activates STING. Nature 2013, doi:10.1038/nature12306.

138. Whiteley AT, Eaglesham JB, de Oliveira Mann CC, Morehouse BR, Lowey B, Nieminen EA, Danilchanka O, King DS, Lee ASY, Mekalanos JJ, et al.: Bacterial cGAS-like enzymes synthesize diverse nucleotide signals. Nature 2019, 567:194–199.

139. Lowey B, Whiteley AT, Keszei AFA, Morehouse BR, Mathews IT, Antine SP, Cabrera VJ, Kashin D, Niemann P, Jain M, et al.: CBASS Immunity Uses CARF-Related Effectors to Sense 3′–5′- and 2′– 5′-Linked Cyclic Oligonucleotide Signals and Protect Bacteria from Phage Infection. Cell 2020, 182:38–49.e17.

140. Cohen D, Melamed S, Millman A, Shulman G, Oppenheimer-Shaanan Y, Kacen A, Doron S, Amitai G, Sorek R: Cyclic GMP–AMP signalling protects bacteria against viral infection. Nature 2019, 574:691–695.

141. Kazlauskiene M, Kostiuk G, Venclovas Č, Tamulaitis G, Siksnys V: A cyclic oligonucleotide signaling pathway in type III CRISPR-Cas systems. Science (80- ) 2017, doi:10.1126/science.aao0100.

142. Niewoehner O, Garcia-Doval C, Rostøl JT, Berk C, Schwede F, Bigler L, Hall J, Marraffini LA, Jinek M: Type III CRISPR-Cas systems produce cyclic oligoadenylate second messengers. Nature 2017, 548:543–548.

143. Kolesnik M V., Fedorova I, Karneyeva KA, Artamonova DN, Severinov K V.: Type III CRISPR-Cas Systems: Deciphering the Most Complex Prokaryotic Immune System. Biochem 2021, 86:1301– 1314.

144. Rivera-Serrano EE, Gizzi AS, Arnold JJ, Grove TL, Almo SC, Cameron CE: Viperin Reveals Its True Function. Annu Rev Virol 2020, doi:10.1146/annurev-virology-011720-095930.

145. Bernheim A, Millman A, Ofir G, Meitav G, Avraham C, Shomar H, Rosenberg MM, Tal N, Melamed S, Amitai G, et al.: Prokaryotic viperins produce diverse antiviral molecules. Nature 2020, in press.

146. Liu Y, Makarova KS, Huang W-C, Wolf YI, Nikolskaya AN, Zhang X, Cai M, Zhang C-J, Xu W, Luo Z, et al.: Expanded diversity of Asgard archaea and their relationships with eukaryotes. Nature 2021, 593:553–557.

147. Zaremba-Niedzwiedzka K, Caceres EF, Saw JH, Bäckström Di, Juzokaite L, Vancaester E, Seitz KW, Anantharaman K, Starnawski P, Kjeldsen KU, et al.: Asgard archaea illuminate the origin of eukaryotic cellular complexity. Nature 2017, 541:353–358.

148. Leão P, Little ME, Appler KE, Sahaya D, Aguilar-Pine E, Currie K, Finkelstein IJ, De Anda V, Baker BJ: Asgard archaea defense systems and their roles in the origin of immunity in eukaryotes. bioRxiv 2023, 10.1038/s41467-024-50195-2.

149. Jean C, Ernest M, Veronica Hernandez Trejo, Florian Tesson, Gal Ofir, Enzo Z. Poirier, Aude Bernheim: Conservation of antiviral systems across domains of life reveals novel immune mechanisms in humans. *bioRxiv* 2023,

150. Jenson JM, Li T, Du F, Ea CK, Chen ZJ: Ubiquitin-like conjugation by bacterial cGAS enhances anti-phage defence. Nature 2023, doi:10.1038/s41586-023-05862-7.

151. Hör J, Wolf SG, Sorek R: Bacteria conjugate ubiquitin-like proteins to interfere with phage assembly. bioRxiv 2023,

152. Bingham R, Ekunwe SIN, Falk S, Snyder L, Kleanthous C: The major head protein of bacteriophage T4 binds specifically to elongation factor Tu. J Biol Chem 2000, doi:10.1074/jbc.M002546200.

153. Depardieu F, Didier JP, Bernheim A, Sherlock A, Molina H, Duclos B, Bikard D: A Eukaryotic-like Serine/Threonine Kinase Protects Staphylococci against Phages. Cell Host Microbe 2016, doi:10.1016/j.chom.2016.08.010.

154. Roy CR, Cherfils J: Structure and function of Fic proteins. Nat Rev Microbiol 2015, doi:10.1038/nrmicro3520.

155. Sberro H, Leavitt A, Kiro R, Koh E, Peleg Y, Qimron U, Sorek R: Discovery of Functional Toxin/Antitoxin Systems in Bacteria by Shotgun Cloning. Mol Cell 2013, doi:10.1016/j.molcel.2013.02.002.

156. Garvey P, Fitzgerald GF, Hill C: Cloning and DNA sequence analysis of two abortive infection phage resistance determinants from the lactococcal plasmid pNP40. Appl Environ Microbiol 1995, doi:10.1128/aem.61.12.4321-4328.1995.

157. Dy RL, Przybilski R, Semeijn K, Salmond GPC, Fineran PC: A widespread bacteriophage abortive infection system functions through a Type IV toxin-antitoxin mechanism. Nucleic Acids Res 2014, doi:10.1093/nar/gkt1419.

158. Dedrick RM, Jacobs-Sera D, Guerrero Bustamante CA, Garlena RA, Mavrich TN, Pope WH, Cervantes Reyes JC, Russell DA, Adair T, Alvey R, et al.: Prophage-mediated defence against viral attack and viral counter-defence. Nat Microbiol 2017, doi:10.1038/nmicrobiol.2016.251.

159. Li Y, Liu X, Tang K, Wang W, Guo Y, Wang X: Prophage encoding toxin/antitoxin system PfiT/PfiA inhibits Pf4 production in Pseudomonas aeruginosa. Microb Biotechnol 2020, doi:10.1111/1751-7915.13570.

160. Alam A, Bröms JE, Kumar R, Sjöstedt A: The Role of ClpB in Bacterial Stress Responses and Virulence. Front Mol Biosci 2021, doi:10.3389/fmolb.2021.668910.

161. Rinke C, Chuvochina M, Mussig AJ, Chaumeil PA, Davín AA, Waite DW, Whitman WB, Parks DH, Hugenholtz P: A standardized archaeal taxonomy for the Genome Taxonomy Database. Nat Microbiol 2021, 6:946–959.

162. Spang A, Caceres EF, Ettema TJG: Genomic exploration of the diversity, ecology, and evolution of the archaeal domain of life. Science (80- ) 2017, 357.

163. Camarillo-Guerrero LF, Almeida A, Rangel-Pineros G, Finn RD, Lawley TD: Massive expansion of human gut bacteriophage diversity. Cell 2021, 184:1098–1109.e9.

