## Supplementary Figure for "Genomic landscape of antiviral defense systems in prokaryotes and phages"

**Highly diverse antiviral defense systems encoded by prokaryote and phage**

**a**

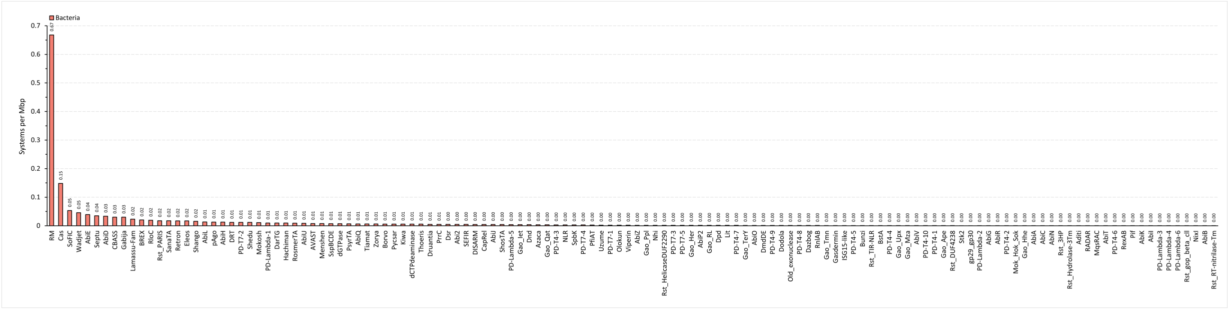


**b**

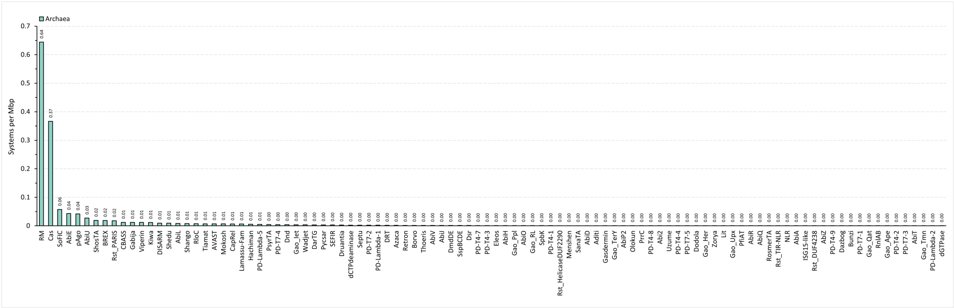


**c**

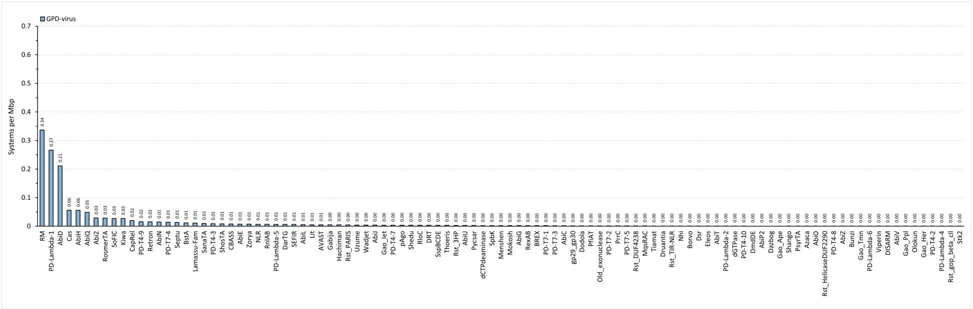


**Supplementary Figure 1. The occurrence frequency of defense systems among bacteria, archaea, and virus domains. a**, Number of defense systems per Mbp in bacterial genomes. **b**, Number of defense systems per Mbp in archaeal genomes. **c**, Number of defense systems per Mbp in viral genomes. Defense systems in the x-axis are listed in descending order of number. Frequency is retained to two decimal places.

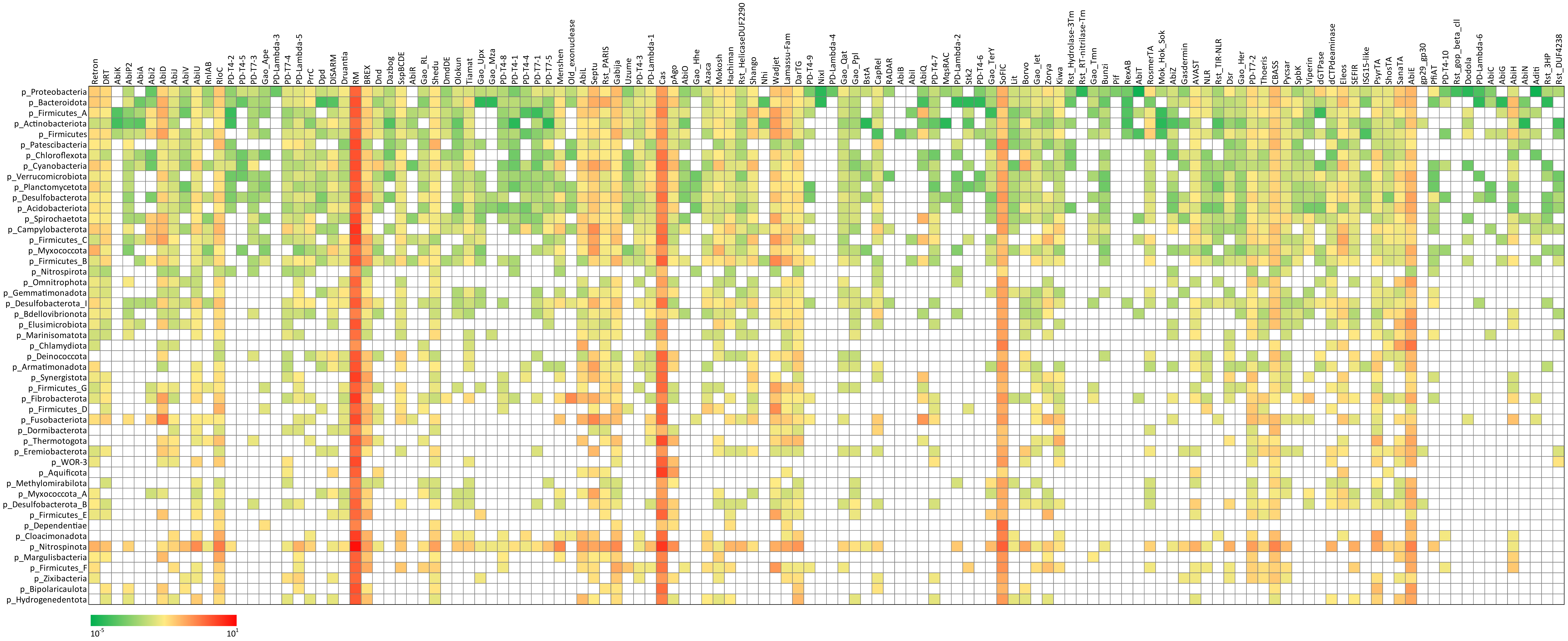


**Supplementary Figure 2. Distribution of defense systems among 49 different bacterial taxa.** Data are shown by the number of systems per Mbp and demonstrated in logarithmic scale colormap.

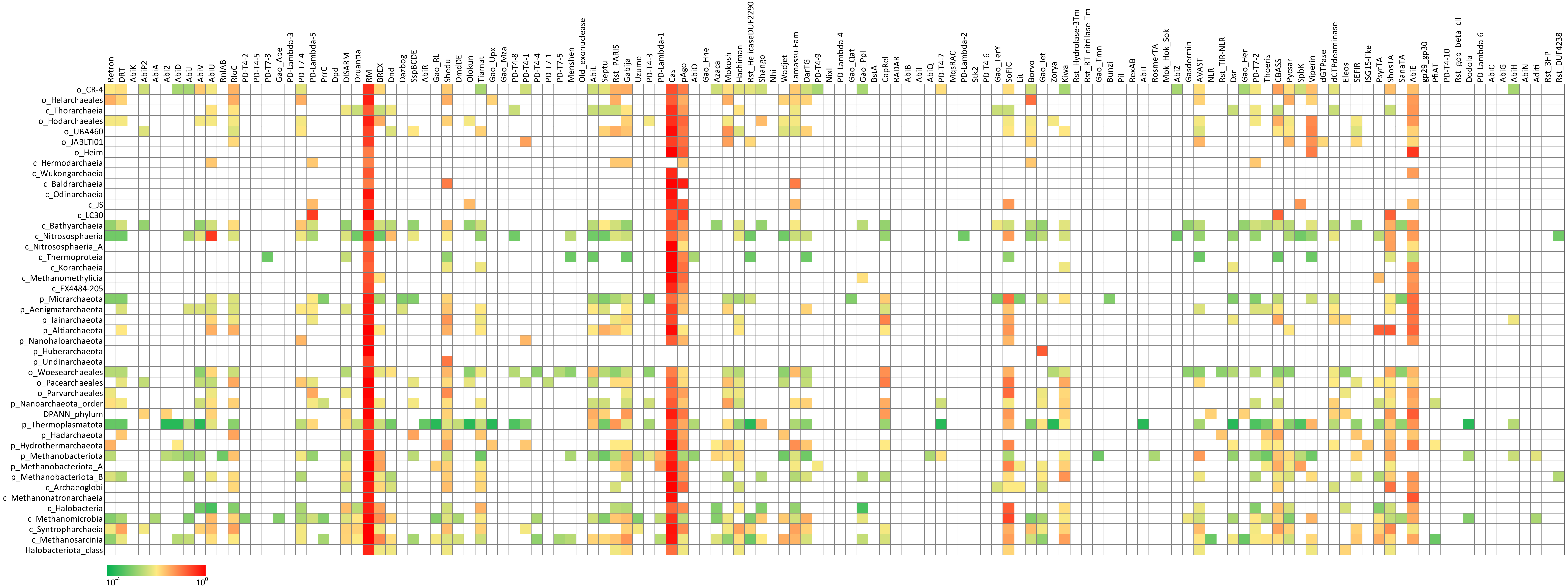


**Supplementary Figure 3. Distribution of defense systems among 45 different archaeal taxa.** Data are shown by the number of systems per Mbp and demonstrated in logarithmic scale colormap.
